# GATA6 Mediates Endoderm and Mesoderm Progenitor Fate Driven by WNT and NODAL Signaling

**DOI:** 10.1101/2025.06.20.660704

**Authors:** Miriam Gordillo, Rebecca Wu, Kelly M. Banks, Neranjan de Silva, Todd Evans

## Abstract

During gastrulation, NODAL and WNT signaling drives distinct gene regulatory networks (GRNs) to promote pluripotency exit, specify progenitor fate, and direct lineage differentiation. Here, we employ murine embryonic stem cell directed differentiation toward endoderm and mesoderm lineages to reveal a crucial role of GATA6 during anterior primitive streak-derived progenitor segregation. GATA6 is known to direct by default extra-embryonic endoderm. To promote definitive endoderm fate, GATA6 reinforces the endoderm GRN driven by NODAL while repressing the mesoderm GRN through down-regulation of WNT activity. During progression of mesoderm differentiation, GATA6 induces a switch in the transcriptional output regulated by WNT, promoting lateral mesoderm specification and repression of paraxial mesoderm fate. Thus, this single transcription factor mediates formation of four distinct tissue types, depending on integration of NODAL/WNT activities. Regulation of *Eomes* and *T/Bra* expression during anterior primitive streak progression is an essential function of GATA6 during the specification of endoderm or mesoderm fate.

## INTRODUCTION

The interaction between signaling pathways and transcriptional networks controls the specification of cell lineages during gastrulation. Definitive endoderm (DE) and mesoderm (ME) development requires the concerted activity of the NODAL and WNT pathways within signaling gradients formed in the primitive streak (Robertson, 2014). In this context, NODAL (a member of the TGFβ superfamily) and WNT cooperate to induce a progenitor population known as mesendoderm (Estarás et al., 2015; Funa et al., 2015; Kubo et al., 2004; Reid et al., 2012; Rodaway and Patient, 2001; Zorn and Wells, 2007). After mesendoderm formation, NODAL and WNT control the segregation of DE and ME progenitors through unknown molecular mechanisms. In vivo, study of the segregation of these lineages is complicated by the transient and rapid mesoderm/endoderm bifurcation in the primitive streak (Mittnenzweig et al., 2021). Work in multiple species indicates that high levels of NODAL promote DE fate, and lower activity levels promote ME (Robertson, 2014). Furthermore, after an initial requirement for WNT signaling, early DE progenitors committed to a SOX17+ DE fate become WNT independent by an unknown mechanism (Pour et al., 2022).

Recent evidence demonstrates the dynamic nature of the interaction of NODAL and WNT to induce different fates. In mice, constitutively active WNT signaling pushes epiblast cells to posterior and middle streak ME progenitors at the expense of anterior ME progenitors, and it was proposed that WNT cooperates with NODAL to promote the differentiation of anterior ME derivatives (Hernández-Martínez et al., 2024). Using geometrically constrained models of human pluripotent cell differentiation, WNT3a was shown to induce NODAL expression, which then drives DE differentiation. In this context, NODAL activity also restricts posterior ME specification, and if NODAL is inhibited, cell fate is diverted to paraxial ME, indicating the ability of NODAL to modulate WNT signaling (Ortiz-Salazar et al., 2024). A similar in vitro approach using CHIR99021 (CHIR), a GSK3β inhibitor that stabilizes β-catenin and is commonly used as a WNT agonist (Bennett et al., 2002), found that small variations in CHIR concentration induced distinct *Nodal* activity levels and dynamics which in turn determined the choice of DE or ME fate (Robles-Garcia et al., 2024).

Interpretation of extracellular signals by transcription factors (TFs) involves cell-type specific protein complexes that drive tissue-specific gene regulatory networks (GRNs). During exit of pluripotency, NODAL and WNT regulate the expression of genes encoding TFs including *T/Bra*, *Eomes*, *Gata4*, and *Gata6*, that dismantle the pluripotency network and establish the GRNs that will drive ME and DE fate (Brown et al., 2011; Dunn et al., 2004; Faial et al., 2015; Funa et al., 2015; Liu et al., 2011; Morgani and Hadjantonakis, 2020; Senft et al., 2018; Zorn and Wells, 2007). BRACHYURY and EOMES are essential regulators of pluripotency exit, neuroectoderm fate repression, and DE and ME differentiation and play pivotal roles during entry and exit from the mesendoderm stage (Arnold et al., 2008; Faial et al., 2015; Gentsch et al., 2017; Herrmann et al., 1990; Tosic et al., 2019). *T/Bra* is a WNT target and direct activator of the ME regulatory network driven by *Tbx6* that controls paraxial ME formation and patterning (Hofmann et al., 2004; Mariani et al., 2021; Yamaguchi et al., 1999). *Eomes* is a critical regulator of the early DE and anterior ME GRNs (Probst and Arnold, 2017). GATA6 is quite remarkable, as it is indispensable for the development of extraembryonic endoderm, while also playing an essential role in DE and ME development and organogenesis (Carrasco et al., 2012; Hayashi et al., 2016; Jun et al., 2023; Keijzer et al., 2001; Kozhemyakina et al., 2014; Morrisey et al., 1998a; Sam et al., 2020, 2024; Xuan et al., 2012; Zhao et al., 2008). Mutations in *GATA6* are associated with human pancreatic and gall bladder agenesis, congenital heart defects, and hepatobiliary malformations (Allen et al., 2012; Chao et al., 2015; De Franco et al., 2013). GATA6 function in DE and ME has recently been shown to involve interactions with EOMES and SMAD2/3 (Bisson et al., 2024; Chia et al., 2019; Heslop et al., 2021, 2022). Despite advances in understanding that NODAL, WNT, and the above TFs regulate the exit of pluripotency and entry into mesendoderm fate, mechanisms at the exit from mesendoderm towards specification of DE or ME remain unclear.

Here, we investigate the role of GATA factors in the transcriptional networks driven by NODAL and WNT during lineage segregation from mesendoderm. Previous reports on the function of *Gata4* or *Gata6* during mouse embryonic stem cell (ESC) differentiation demonstrated their capacity to direct visceral endoderm but not DE development (Capo-chichi et al., 2010; Fujikura et al., 2002; Holtzinger et al., 2010; Morrisey et al., 1998b; Soudais et al., 1995). Using serum-free conditions to avoid the known inhibitory effect of FBS on DE differentiation (D’Amour et al., 2005; Kubo et al., 2004), we confirmed a critical role for *Gata6* in murine DE differentiation, consistent with results using human ESCs (Chia et al., 2019; Fisher et al., 2017; Shi et al., 2017; Tiyaboonchai et al., 2017). We find that *Gata6* expression in mesendoderm progenitors drives DE fate by reinforcing and propagating the DE GRN controlled by NODAL while repressing the ME GRN controlled by WNT. Yet, unexpectedly, during ME differentiation induced by sustained WNT activation, GATA6 promotes lateral plate ME fate at the expense of paraxial ME by modulating the transcriptional output of WNT. *Eomes* and T/*Bra* are among the first targets of GATA6-mediated transcriptional regulation, essential for the transition of mesendoderm progenitors to DE or ME. In this manner, a single TF, GATA6, directs early steps of organogenesis from distinct germ layers for specification of four distinct early progenitor fates.

## RESULTS

### Expression of GATA Factors Drives Definitive Endoderm Specification

We used a defined serum-free medium and Activin A to efficiently induce DE progenitors from mouse ESCs (Figure S1A). To track DE progenitors, flow cytometry was used to analyze the c-KIT and CXCR4 double-positive population (Gouon-Evans et al., 2006). However, in no-Activin or low-Activin conditions, these markers were unreliable compared to the combination of EpCAM/CXCR4 to discriminate DE from other populations (Figure S1B, S1C). Thus, although both combinations were analyzed as validation, we quantified the EpCAM/CXCR4 population as DE throughout this study. High concentrations of Activin A promote specification of DE, while concentrations below 25 ng/ml do not induce DE progenitors or the expression of DE markers (Figure S1C and S1D). Transcriptome analysis shows enrichment of DE signatures derived from single-cell RNA-seq of mouse embryos with 75 ng/ml Activin A (Figure S1E). In a time-course analysis for the expression of DE TFs induced by Activin A at 75 ng/ml, we observed three waves of expression of transcription factors of the DE Gene Regulatory Network (GRN, Figure S1F). *Eomes*, *FoxA2*, and *Mixl1* are the earliest expressed followed by *Gata6*, *Sox17*, *FoxH1* and *Lhx1*, and finally *Gata4*, *Hesx1*, *Hhex*, and *FoxA3* (blue, green, and red lines, respectively, Figure S1F).

To investigate the role of *Gata4* and *Gata6* in DE development, we first tested the ability of each to drive DE development using previously established mESC lines that allow conditional expression of *Gata4* or *Gata6* upon the addition of doxycycline (+Dox; Figure 1A) (Holtzinger et al., 2010; Turbendian et al., 2013). Initial induction of Dox under conditions of 75 ng/ml Activin showed a significant increase in DE progenitors, suggesting that forced expression of GATA4 or GATA6 can enhance the effect of Activin A during DE specification (Figure 1B, GATA6; Figure S1G, GATA4). Next, the GATA factors were induced using just 10 ng/ml Activin A, a concentration that on its own does not specify DE (Figure S1C). Indeed, DOX-induction of either GATA factor generates a significant DE population comparable in numbers to that induced by 75 ng/ml Activin A (Figure 1B, *Gata6*; Figure S1G, *Gata4*). qPCR analysis confirmed the increased expression of multiple DE markers (Figure 1C, *Gata6*; Figure S1H, *Gata4*). Gene list analysis of the *Gata6*-induced transcriptome using Enrichr (Chen et al., 2013) shows a high enrichment of genes associated with DE development (Figure S1I). Gene Set Enrichment Analysis (GSEA) using sets extracted from embryonic mouse single cell RNA-seq data in the literature shows enrichment of early DE signatures (Figure 1D).

**Figure 1.**
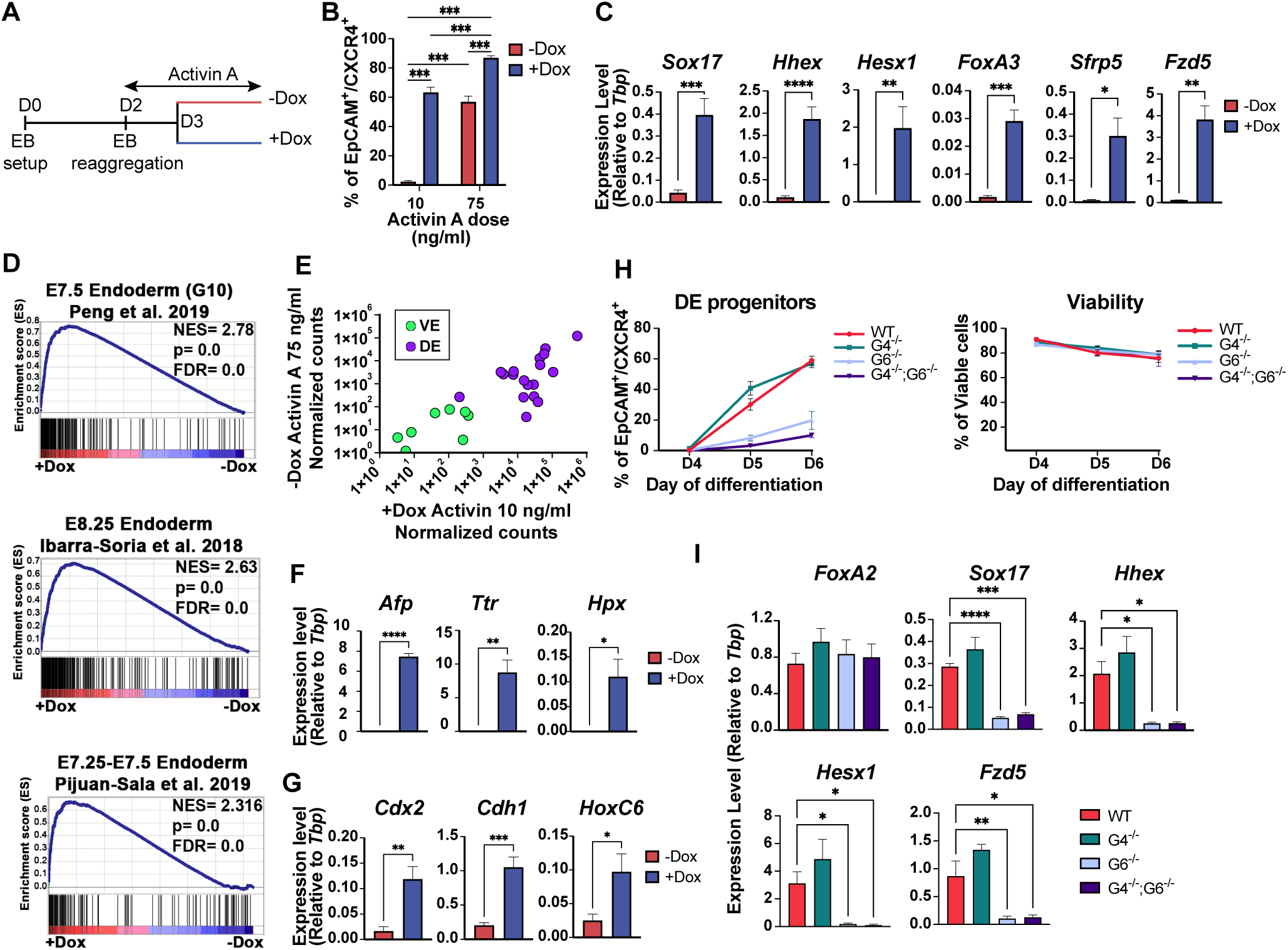
Expression of GATA4 or GATA6 can drive murine definitive endoderm specification, and GATA6 is required. (A) Schematic diagram of the protocol for directed differentiation and induced expression of GATA factors. Embryoid body (EB) formation was initiated on day 0 and cells reaggregated on day 2 in the presence of Activin A. Doxycycline (+Dox, 30 ng/ml) was added (or not, -Dox) to induce GATA factor expression starting on day 3. (B) Quantification by flow cytometry of CXCR4+/EpCAM+ DE progenitors on day 4.5 of differentiation after induction of *Gata6* expression. See Figure S1B for gating strategy. (C) Gene expression analysis by RT-qPCR for key DE markers expressed on day 4 of differentiation using 10 ng/ml Activin A. (D) GSEA analysis of RNA-seq gene expression data at day 4 of differentiation (+Dox *vs* –Dox; 10ng/ml Activin A) using mouse embryo endoderm enriched gene sets obtained from single-cell RNA-seq data previously published (Ibarra-Soria et al., 2018; Peng et al., 2019; Pijuan-Sala et al., 2019). NES=normalized enrichment score; FDR=false discovery rate. (E) Gene expression of DE (purple) and VE (green) classifier gene sets previously identified through single-cell RNA-seq of mouse embryo cells (Nowotschin et al., 2019). Expression as normalized counts was obtained from RNA-seq data at day 4 of differentiation of cells grown in 10 ng/ml Activin A, +Dox, compared to cells grown in 75 ng/ml Activin A. (F and G) RT-qPCR analysis of tissue markers in DE progenitors that underwent further hepatic differentiation (F) or mid-hindgut differentiation (G). (H) DE differentiation of WT, *Gata4*^-/-^, *Gata6*^-/-^, and *Gata4*^-/-^*/Gata6*^-/-^ mESCs. The left panel shows quantification of DE progenitors by flow cytometry; right panel shows viability analysis by flow cytometry. Cells were differentiated as depicted in Figure S1A in the presence of 75 ng/ml Activin A. (I) Gene expression analysis by RT-qPCR of key DE markers in WT, *Gata4*^-/-^, *Gata6*^-/-^, and *Gata4*^-/-^*/Gata6*^-/-^ mESCs. Data in B, C, F, G, and I are presented as mean ± SEM. *n* ≥ 3 biologically independent experiments. Statistical significance in C, F, and G was evaluated by unpaired, two-tailed Student’s t test for two-group comparisons. Statistical significance in B was evaluated with two-way ANOVA followed by Tukey’s test for multiple comparisons. Statistical significance in I was evaluated with one-way ANOVA followed by Dunnett’s test for multiple comparisons relative to WT.

The early DE lining of the primitive gut tube comprises a mixture of DE and VE progenitors with highly similar overall gene expression profiles (Nowotschin et al., 2019). However, DE and VE cells can be distinguished using specific classifier gene sets identified through single-cell RNA-seq of embryo cells. Using these classifier sets, we confirmed that the DE progenitors generated by induction of GATA factors correspond to DE progenitors that express low levels of genes predictive of the VE class and high levels of genes predictive of the DE class (Figure 1E). In addition, these sets of genes are expressed at a similar level as the cells differentiated with a high concentration of Activin, indicating that GATA factors direct the same type of DE generated by high activity of the NODAL/Activin signaling pathway. Replating Day 5 EBs in hepatoblast or mid-hindgut differentiation conditions showed that DE progenitors generated by expression of GATA factors differentiate efficiently into these cell types derived from anterior or posterior DE (Figure 1F-G, *Gata6*; Figure S1J-K, *Gata4*).

Using mouse *Gata6^-/-^*; *Gata4^-/-^*; and *Gata4^-/-^*,*Gata6^-/-^* ESCs generated by gene targeting (Molkentin et al., 1997; Zhao et al., 2008, 2005), *Gata6* (and not *Gata4*) function was found to be essential for DE formation (Figure 1H-I). An essential role for GATA6 in DE development was also reported using human pluripotent cells (Fisher et al., 2017; Shi et al., 2017; Tiyaboonchai et al., 2017), and our data indicate it is a conserved program. However, in contrast to human pluripotent cells, the deficiency in DE development from murine *Gata6* null cells is not the result of cell death (Figure 1H) (Fisher et al., 2017; Tiyaboonchai et al., 2017). Taken together, our results demonstrate the capacity of either GATA4 or GATA6 to induce functional DE progenitors and show a conserved requirement for *Gata6* in DE formation.

### Gata6 Reinforces the NODAL Pathway to Drive Endoderm Specification

After induction of *Gata6* in the context of a low dose of Activin A, expression levels of *Nodal* and its direct downstream target, *Lefty1*, are increased (Figure 2A). In addition, enrichment analysis of the GATA6-induced transcriptome highlights gene sets regulated by the TGFβ/NODAL signaling pathway and SMAD2, SMAD3, and SMAD4, the main effectors of the NODAL pathway (Figure S2A-S2B). GSEA using sets of genes regulated by SMADs gathered from previous studies (Chia et al., 2019; Chiu et al., 2014; Heslop et al., 2022; Kim et al., 2011; Senft et al., 2018) also reveals the up-regulation of multiple sets of direct SMAD2/3 targets after Dox-induction of *Gata6* (Figure 2B). qPCR analysis of SMAD2/3/4 direct targets *Lhx1*, *Cer1*, and *Eomes* (Chiu et al., 2014), shows that transcript levels are Activin A dose-dependent (Figure S2C) and that *Gata6* (Figure 2D) or *Gata4* (Figure S2D) upregulate the expression of these genes. RNAseq analysis shows the extensive transcriptional regulation of other multiple SMAD2/3/4 direct targets by GATA6 (Figure 2C). Transcript levels of *FoxH1*, a TF essential for NODAL signaling and DE specification (Chiu et al., 2014; Hoodless et al., 2001; Slagle et al., 2011), are also Activin A dose-dependent (Figure S2C) and enhanced by GATA factors (Figure 2D, *Gata6*; Figure S2D, *Gata4*). Furthermore, direct targets of FOXH1 are also upregulated by expression of *Gata6*, as indicated by GSEA analysis (Figure 2B). Expression analysis of SMAD2/3 direct targets *Eomes*, *Lhx*, and *Cer1* in *Gata6^-/-^* cells shows that *Gata6* is necessary to maintain their expression (Figure 2E). However, there is no significantly impaired expression level of *FoxH1* or *Nodal* or decreased pathway activity as reflected by the normal expression level of *Lefty1* in the absence of GATA6 (Figure S2E). To test the ability of GATA6 to specify DE independently of NODAL signaling, *Gata6* expression was induced with Dox in the absence of Activin A, and this failed to generate DE (Figure 2F). In addition, inhibition of Activin type I receptors ALK5, ALK4, and ALK7 with the A83-01 inhibitor at low or high doses of Activin A blocked DE specification despite the forced expression of *Gata6* (Figure 2G and S2F), indicating that *Gata6* requires a NODAL signal to induce DE. Together, these results indicate that *Gata6* is an essential component of the NODAL regulated network that reinforces and propagates activity of the pathway to drive the DE regulatory network.

**Figure 2.**
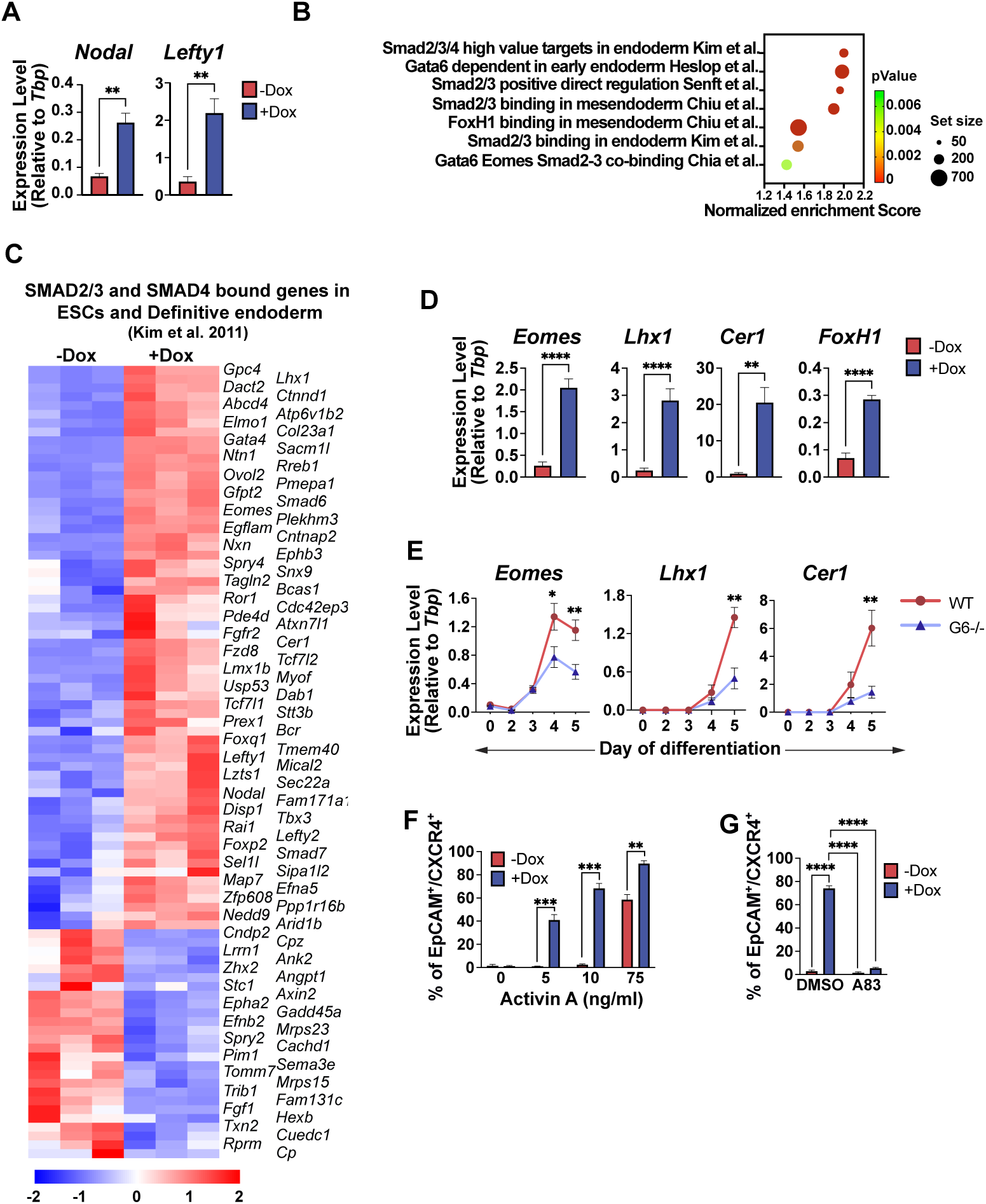
GATA6 reinforces the NODAL pathway. (A) Gene expression analysis by RT-qPCR for NODAL pathway markers *Nodal* and *Lefty1* on day 4 of differentiation using 10 ng/ml Activin A). (B) GSEA using the *Gata6*-induced transcriptome comparing sets of genes regulated by TGFβ/NODAL pathway effectors, as described from previous studies, including direct SMAD2/3 targets. (C) The heat map from *Gata6*-induced RNA-seq analysis shows the extensive transcriptional regulation of multiple SMAD2/3/4 direct targets as defined previously (Kim et al., 2011). (D) *Gata6*-induced gene expression analysis by RT-qPCR for known SMAD2/3/4 direct targets on day 4 of differentiation using 10 ng/ml Activin A. (E) Gene expression analysis by RT-qPCR for *Eomes*, *Lhx1*, and *Cer1* during endoderm differentiation using 75 ng/ml Activin A shows that *Gata6*^-/-^ cells cannot maintain expression of these genes. (F) Based on flow cytometry, specification of DE is dose dependent, and without induction of *Gata6* expression requires a high dose of Activin A, but with induction of *Gata6* a low dose is sufficient. Nevertheless, with no-dose (0 ng/ml), *Gata6* fails to induce DE. (G) As quantified by flow cytometry, specification of DE by induction of *Gata6* (in 10 ng/ml Activin A) requires active Nodal/Activin A signaling, as it is blocked by addition of the A83-01 ALK4/5/7 type I receptor inhibitor. For all panels, n≥3 independent differentiation experiments. Data in A, D-G are presented as mean ± SEM. Statistical significance in A, D-F was evaluated by unpaired, two-tailed Student’s t test for each two-group comparison. Statistical significance in G was evaluated with two-way ANOVA followed by Tukey’s test for multiple comparisons.

### GATA Factors Promote the Transition of Mesendoderm Toward Definitive Endoderm Fate

We hypothesized that GATA factors drive mesendoderm exit towards DE fate at the expense of ME fate. GSEA analysis of the transcriptome of cells at low concentration of Activin A with or without Dox revealed the capacity of GATA6 (Figure 3A) or GATA4 (Figure S3A) to promote the DE regulatory network and that DE specification is coupled with their ability to suppress the ME regulatory network. The qPCR analysis for transcript levels of key ME TFs and markers at this early stage of differentiation confirms that expression of *Gata6* (Figure 3B) or *Gata4* (Figure S3B) causes down-regulation of genes essential for ME development. In mice, mesendoderm progenitors are identified by the co-expression of *FoxA2* and *T/Bra* (Kubo et al., 2004). After segregation from mesendoderm, DE progenitors upregulate *FoxA2* and downregulate *T/Bra*, while ME progenitors down-regulate the expression of *FoxA2* and upregulate *T/Bra* (Figure S3E). The qPCR analysis 24 hr after Dox-induction of *Gata6* (Figure 3C) or *Gata4* (Figure S3C) shows upregulation of *FoxA2* and down-regulation of T/*Bra*. Flow cytometry analysis confirms the down-regulation of *T/Bra* expression (Figure 3D, *Gata6*; Figure S3D, *Gata4*). Flow cytometry analysis for FOXA2 was not possible using currently available antibodies. Therefore, to monitor co-expression of *FoxA2* and *T/Bra*, we generated a *Gata4* inducible cell line in the background of a dual reporter line containing GFP cDNA targeted to the *T/Bra* locus and a truncated version of the human CD4 cDNA targeted to the *Foxa2* locus (Cheng et al., 2008). This cell line allowed evaluation of the progression of double-positive (GFP-BRA^+^/CD4-FOXA2^+^) mesendoderm progenitors after the induction of *Gata4*. Kinetic analysis of this population shows that Dox-induction of *Gata4* results in a steady increase of the CD4-FOXA2^+^/GFP-BRA^-^ population (Figure S3F), while the mesendoderm GFP-BRA^+^/CD4-FOXA2^+^ population decreases (Figure S3G). Thus, the results indicate that GATA factors promote the exit of mesendoderm progenitor towards DE fate at the expense of ME fate.

**Figure 3.**
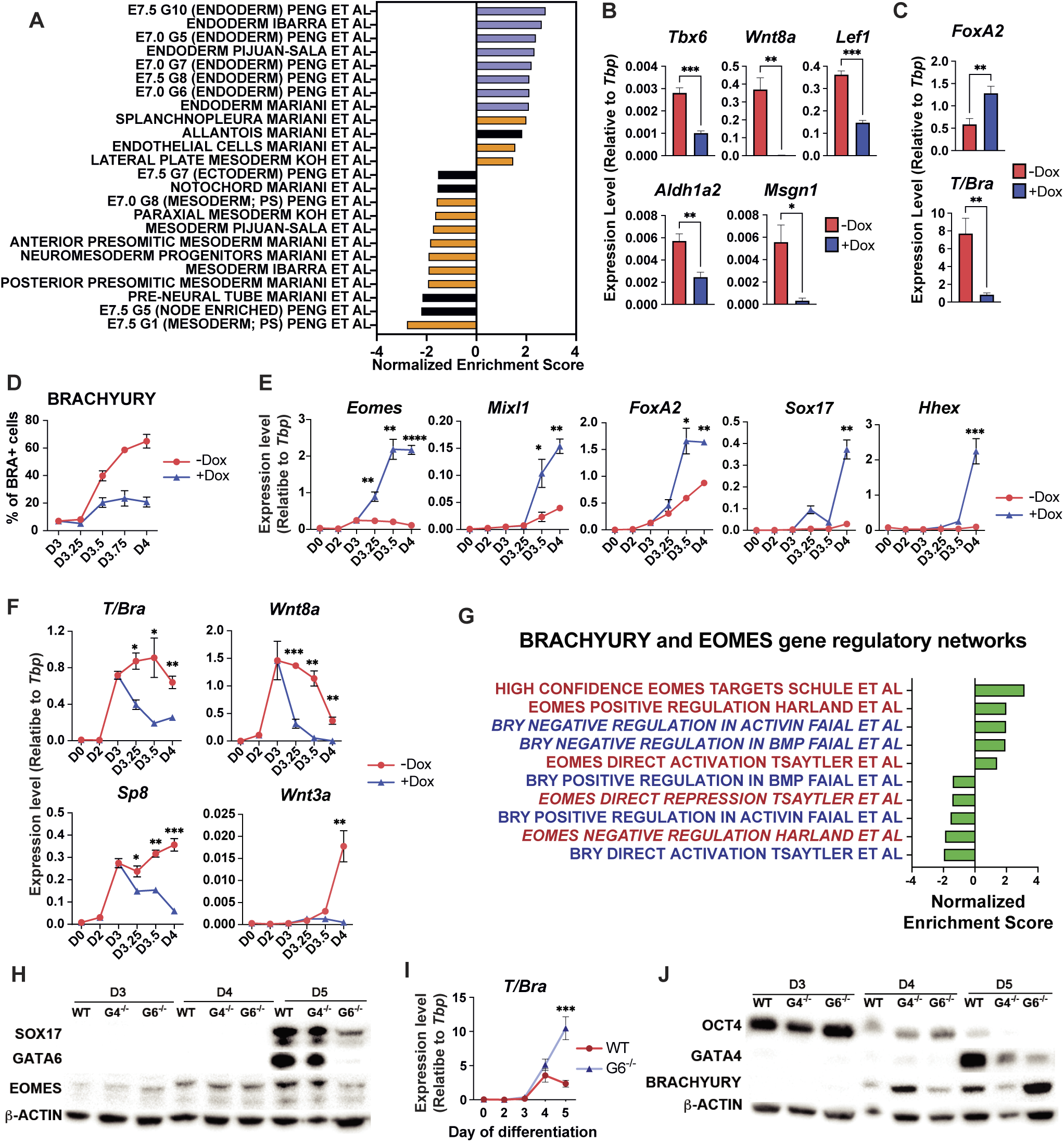
GATA6 drives mesendoderm cells to definitive endoderm fate. (A) GSEA using RNA-seq data of cells at day 4 with or without Dox-induced expression of *Gata6* at low concentration of Activin A shows enrichment for gene sets representing the definitive endoderm regulatory network and suppression for those of the mesendoderm regulatory network. (B) Gene expression analysis by RT-qPCR for key mesoderm drivers on day 4 of differentiation following Dox-induction of *Gata6* and culture in 10 ng/ml Activin A. (C) Gene expression analysis by RT-qPCR for *FoxA2* and *T/Bra* on day 4 of differentiation using 10 ng/ml Activin A. (D) Flow cytometry quantification for BRA+ cells at defined stages of directed differentiation with or without induction of *Gata6* using 10 ng/ml Activin A. (E and F) Gene expression analysis by RT-qPCR with or without Dox-induced expression of *Gata6* for key early drivers of DE specification (E) or ME specification (F) from day 3 to day 4 of differentiation using 10 ng/ml Activin A. (G) GSEA of RNA-seq data using gene sets regulated by EOMES or BRACHYURY, as reported in previous studies. (H) Representative western blotting analysis using lysates from WT, *Gata4*^-/-^ or *Gata6*^-/-^ lines for expression of SOX17 and EOMES shows that both proteins fail to accumulate in cells lacking GATA6 at day 5 of differentiation using 75 ng/ml Activin A. (I) Gene expression analysis by RT-qPCR shows a failure to down-regulate *T/Bra* at day 5 in cells lacking GATA6 during differentiation using 75 ng/ml Activin A. (J) Western blotting analysis using lysates from WT, *Gata4*^-/-^ or *Gata6*^-/-^ lines for expression of OCT4, GATA4, or BRA confirms accumulation of BRA in cells lacking GATA6 at day 5 of differentiation using 75 ng/ml Activin A. Data in B-F and I are presented as mean ± SEM. Statistical significance was evaluated by unpaired, two-tailed Student’s t test for two-group comparisons.

To assess how GATA factors promote the transition to DE while repressing ME fate, the temporal expression levels of key DE and ME transcription factors were analyzed following the Dox-induced expression of *Gata6*. *Eomes* is the earliest DE transcription factor upregulated by 6 hr following *Gata6* induction (Figure 3E). In contrast, *T/Bra* is significantly downregulated as early as six hours after *Gata6* induction (Figure 3F). GSEA analysis of RNAseq data using gene sets regulated by EOMES and BRACHYURY, as reported in previous studies (Faial et al., 2015; Harland et al., 2021; Schüle et al., 2023; Tsaytler et al., 2023) revealed that gene signatures positively regulated by EOMES are generally upregulated upon *Gata6* induction, whereas those positively regulated by BRACHYURY are down-regulated (Figure 3G). Moreover, genes negatively regulated by EOMES showed decreased expression levels following *Gata6* induction, while those negatively regulated by BRACHYURY were upregulated. The expression levels of *Eomes* and T/*Bra* were next measured during directed specification of DE in cells lacking *Gata6*. The qPCR analysis revealed that *Eomes* transcript levels decrease at day 4-5 of Activin-induced DE differentiation in *Gata6^-/-^* cells, and western blotting assays confirmed that EOMES protein fails to accumulate at the same time (Figure 2E and 3H). In contrast, qPCR and western blotting analyses demonstrated that *T/Bra* expression levels are increased and maintained at higher levels in *Gata6^-/-^* cells during differentiation toward DE (Figure 3I and 3J), indicating that *Gata6* is necessary for the downregulation of *T/Bra* during DE specification. Collectively, these results strongly indicate that GATA6 promotes specification into DE fate predominantly through activation of the TGFβ/NODAL endodermal regulatory network via activating expression of *Eomes*. Concurrently, GATA6 inhibits the mesodermal gene regulatory network by suppressing *T/Bra* expression.

### GATA Factors Negatively Regulate WNT Signaling and Grant WNT Independence to Endoderm Progenitors During Endoderm Specification

GSEA analysis and GO term and pathway enrichment analysis of differentially down-regulated genes following Dox-induced *Gata4/6* expression revealed WNT/β-catenin signaling, an essential regulator of ME formation (Arnold and Robertson, 2009), as one of the main pathways down-regulated by the expression of *Gata6* (Figure 4A) or *Gata4* (Figure S4A). As *T/Bra* is a WNT target and WNT signaling in concert with T box factors (including *T/Bra* and *Tbx6*) regulate ME development (Arnold et al., 2000; Hofmann et al., 2004; Mariani et al., 2021), we hypothesized that GATA factors modulate WNT signaling during ME and DE lineage choice. *Axin2* is widely used as a readout for the activity of the canonical WNT pathway (Jho et al., 2002), and we confirmed that *Axin2* transcript levels are reduced following Dox-induced expression of either *Gata6* (Figure 4B) or *Gata4* (Figure S4B). In contrast, transcript levels of *Axin2* were maintained abnormally high in *Gata6^-/-^* cells during the transition to DE and beyond, indicating that *Gata6* function is necessary to regulate WNT signaling activity during DE specification (Figure 4C and S4C). To better understand the activity of the WNT pathway in this in vitro system, the expression level of *Axin*2 at different concentrations of Activin A was evaluated from Day 2, when Activin A was added, to Day 5, when DE specification was complete. Activity of the pathway (based on *Axin2* transcript levels) increases sharply in the first 24 hr but is only maintained at Activin concentrations below 25 ng/ml (Figure 4D). Interestingly, with Activin A at concentrations that can induce DE progenitors (25 ng/ml and above), the peak of activity at day 3 is followed by a sharp decrease in the next 24 hr when transcript levels of *Axin2* are significantly lower compared to lower concentrations of Activin A conditions (Figure 4D and S4D). This suggests that WNT signaling activity is quickly down-regulated during the transition of mesendoderm to DE, which is in agreement with recent results with human ESCs showing that WNT signaling is required initially for DE induction, yet cells already specified to DE fate become independent of external WNT activation (Pour et al., 2022).

**Figure 4.**
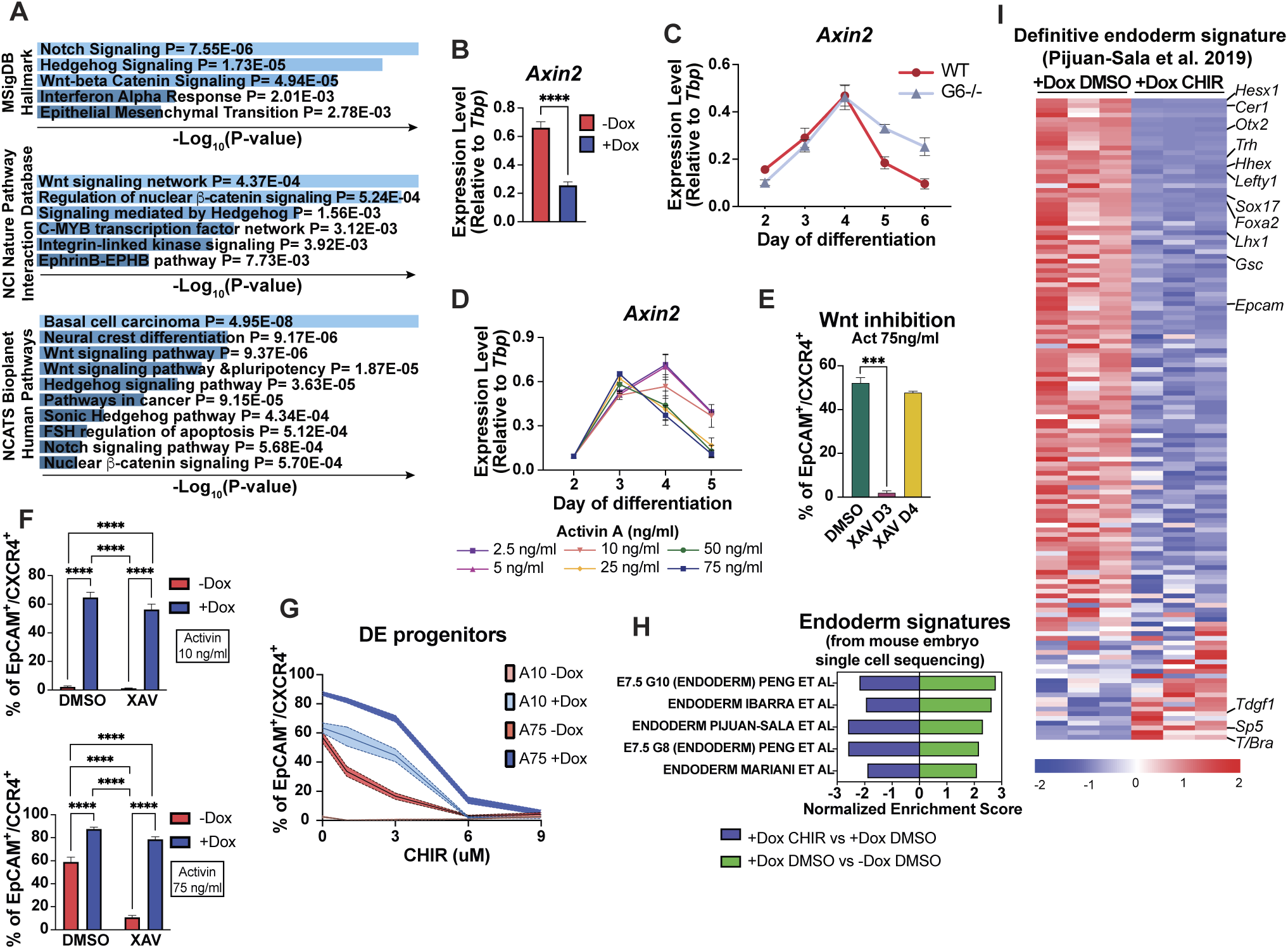
GATA6 negatively regulates WNT signaling and grants WNT independence during DE specification. (A) GO term and pathway enrichment analysis of differentially down-regulated genes at day 4 following Dox-induced *Gata6* expression shows WNT/β-catenin signaling as one of the main down-regulated pathways. (B) Gene expression analysis by RT-qPCR shows down-regulation of the canonical WNT direct target gene *Axin2* at day 4 in cells induced to express *Gata6* during differentiation using 10 ng/ml Activin A. (C) Gene expression analysis by RT-qPCR to measure relative *Axin2* transcript levels at indicated time points in cells lacking GATA6 compared to WT cells during differentiation using 75 ng/ml Activin A. Statistical analysis is reported in Figure S4C. (D) Gene expression analysis by RT-qPCR at indicated time points to measure relative *Axin2* transcript levels in WT cells induced by different concentrations of Activin A. Statistical analysis is reported in Figure S4D. (E) WT cells were induced at day 2 to promote DE specification with 75 ng/ml Activin A and then inhibited for WNT signaling with XAV939 (XAV) at day 3 or day 4. Flow cytometry was used to quantify DE specification compared to control vehicle-treated (DMSO) samples. (F) Analysis for DE specification at day 5 was carried out as in (E) except with (+Dox) or without (-Dox) induction at day 3 of *Gata6* expression in either 10 ng/ml (top) or 75 ng/ml Activin A (bottom). (G) Analysis for DE specification at day 5 was carried out as in (F) except with addition at day 3 of 0, 3, 6, or 9 uM of the WNT pathway activator CHIR99021 (CHIR). (H) GSEA of RNA-seq data at day 4 comparing Dox-induced cells versus DMSO control (green) and Dox-induced cells + 6 μM CHIR versus DMSO control (purple) using endoderm-related gene sets from embryo single-cell RNA-seq experiments. Cells were differentiated with 10 ng/ml Activin A. (I) Heat maps for a similar comparison as in (H) using a definitive endoderm signature gene set (Pijuan-Sala et al., 2019). *n* ≥ 3 biologically independent experiments. Data in B-G are presented as mean ± SEM. Statistical significance in B was evaluated by unpaired, two-tailed Student’s t test for two-group comparisons. Statistical significance in E was evaluated with one-way ANOVA followed by Dunnett’s test for multiple comparisons relative to WT. Statistical significance in F was evaluated with two-way ANOVA followed by Tukey’s test for multiple comparisons.

To confirm the time dependency of WNT signaling in DE specification at 75 ng/ml Activin A, WNT signaling was inhibited on either Day 3 or Day 4 during differentiation before DE progenitors emerge (Figure 1H, WT cells). Inhibition of WNT signaling by adding the XAV939 (XAV) compound on Day 3 but not on Day 4 represses DE specification (Figure 4E). To test the hypothesis that GATA factors repress WNT and thereby program progenitors to be WNT independent, *Gata6* was induced with Dox in cells that were simultaneously inhibited for WNT with XAV at Day 3. Strikingly, induction of *Gata6* expression at either low or high concentrations of Activin A rescues DE specification when WNT is inhibited, indicating that progenitors already expressing *Gata6* no longer depend on WNT signaling (Figure 4F). The qPCR analysis of *Axin2* transcript levels confirmed the inhibitory effect of XAV in both -Dox and +Dox conditions (Figure S4E). A similar rescue of DE fate caused by WNT inhibition is observed with the induction of *Gata4* (Figure S4F).

To test if the ability to repress WNT is necessary for the specification of DE by GATA factors, activation of the WNT pathway was enforced with increasing doses of CHIR. DE progenitor specification using a high dose of Activin A without Dox is inhibited by concentrations of CHIR as low as 1 μM. In contrast, with Dox-induction of *Gata6* (Figure 4G and S5D), at least 6 μM CHIR is required to inhibit DE specification significantly. The qPCR analysis of *Axin2* levels indicates that cells respond to the activation of the pathway but require a higher concentration of CHIR to achieve the same level of *Axin2* expression compared to cells not induced by Dox (Figure S4G). RNAseq analysis confirmed down-regulation of the DE gene regulatory network directed by GATA6 after adding 6 μM CHIR (Figure 4H and 4I). Taken together, these results show that GATA6 negatively regulates WNT signaling as progenitors exit the mesendoderm state towards DE fate, and GATA6 expression renders these progenitors WNT independent during commitment to the DE lineage.

### Gata6 Regulation of WNT Signaling Transcriptional Output is a Key for Mesoderm Subtype differentiation

Like DE, ME development requires signaling from multiple pathways, including NODAL and WNT/β-Catenin (Salehin et al., 2022; Stemple, 2001). Because activation of WNT signaling, one of the main drivers of ME development (Cadigan and Nusse, 1997; Dunty et al., 2008; Galceran et al., 1999), repressed DE specification (Figure 4G), we tested if DE fate was being changed to ME by the activation of WNT. In contrast to DE, there is no single marker or combination of markers to distinguish all ME progenitors. Thus, we quantified the expression of DLL1, FLK1, and PDGFRα. DLL1 is a marker of paraxial ME (Bettenhausen et al., 1995); FLK1 is a marker of lateral plate, cardiovascular, and hematopoietic ME (Kataoka et al., 1997; Yamaguchi et al., 1993); and PDGFRα is a marker for a broad range of ME types, including early paraxial and lateral plate, cardiovascular progenitors, and skeletal ME progenitors (Kataoka et al., 1997; Loh et al., 2016; Orr-Urtreger et al., 1992). Using these markers, we defined two ME subtypes: DLL1-positive ME (DLL1+/PDGFRα+ and DLL1+/PDGFRα-) or DLL1-negative (non-DLL) ME (PDGFRα+/FLK1+, PDGFRα+/FLK1- and PDGFRα-/FLK1+). In all our experiments, no significant amount of DLL1+/FLK1+ cells were found.

Using Activin A alone, we found that 15-20% of DLL1+ or PDGFRα+ ME cells are produced at low or high concentrations of Activin A, respectively, reflecting different subtypes induced in the primitive streak (Figure S5A). Results for each subtype in all different conditions in which Activin A, WNT, and *Gata6* are manipulated are found in Figure S5C. After adding CHIR in -Dox conditions, low doses of Activin A efficiently induce DLL1+ progenitors, whereas high concentrations of Activin A are required to generate non-DLL1 progenitors (Figures 5A and 5B). Additionally, DLL1+ progenitors are generated in a CHIR dose-dependent manner, whereas non-DLL1 progenitors are induced by 1 or 3 μM CHIR but not by 6 or 9 μM. Importantly, in cultures without Dox, endogenous *Gata6* expression level is associated with concentrations of Activin and CHIR that generate a high yield of DE or non-DLL1 progenitors (Figure S5B). Strikingly, after Dox-induction of *Gata6*, DLL1+ ME is largely abrogated, with only a small percentage generated at 9 μM CHIR in low-Activin A conditions (Figure 5A, 5B, and S5C). Furthermore, at high concentrations of Activin A, Dox-induction of *Gata6* completely blocks the generation of DLL1+ ME and results in progenitors that are FLK1+ (positive or negative for PDGFRα) (Figure 5B and S5C). The qPCR analysis further confirmed that WNT activation (using CHIR) in the absence of *Gata6* expression (-Dox conditions) strongly down-regulates DE markers and upregulates paraxial ME markers (Figure S5D). In contrast, dose-dependent WNT activation and Dox-induced *Gata6* expression strongly increase levels for markers of lateral plate ME while repressing paraxial ME marker expression levels, indicating that ME progenitors directed by *Gata6* resemble lateral plate ME at the expense of paraxial ME (Figure S5D).

**Figure 5.**
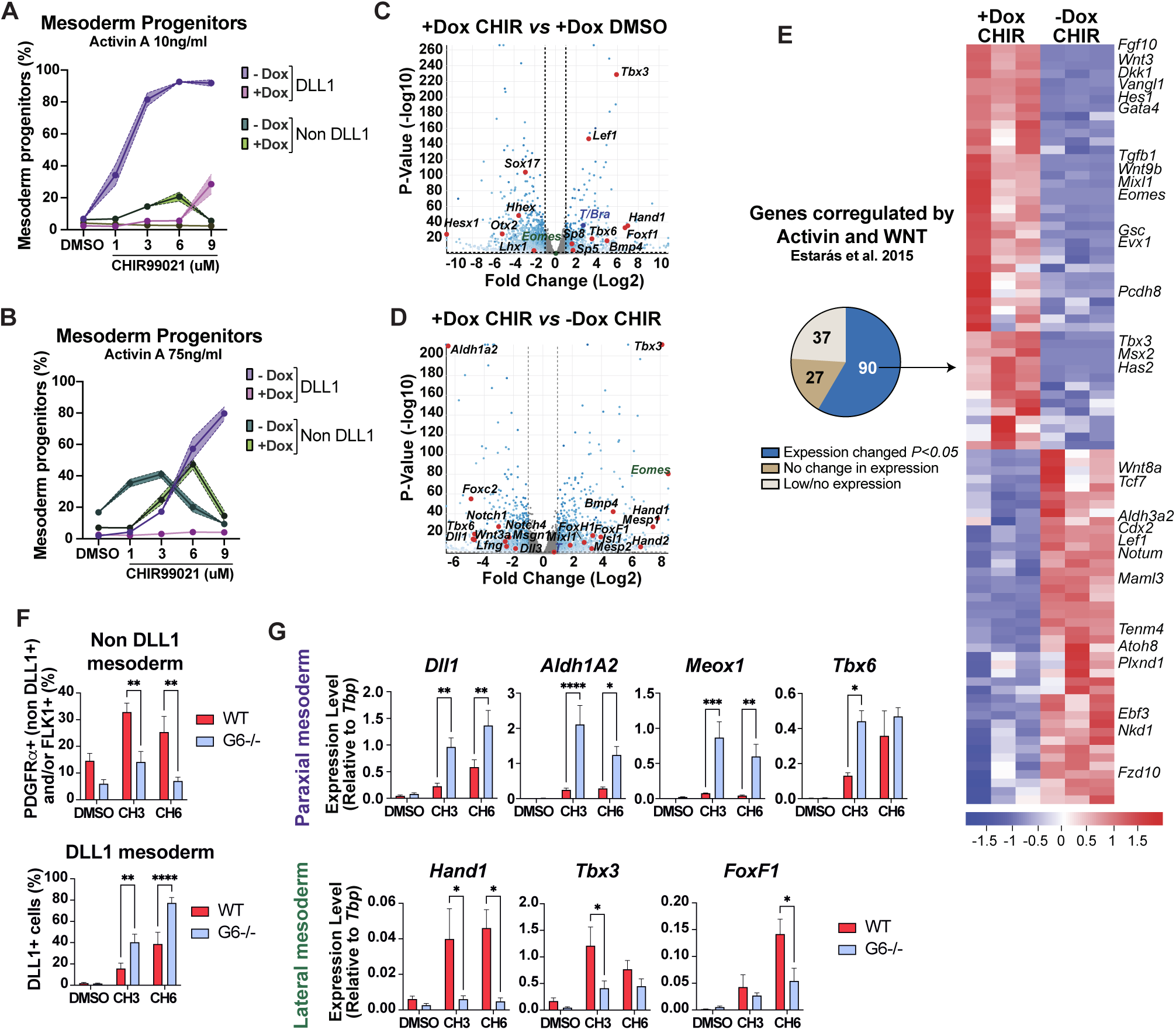
GATA6 cooperates with WNT signaling to specify mesoderm sub-type. (A) Flow cytometry using antibodies to quantify DLL1 subtype mesoderm (DLL1+ regardless of PDGFRα expression) or non-DLL subtype of mesoderm (combining PDGFRα+/FLK1+, PDGFRα+/FLK1-, and PDGFRα-/FLK+) with cells induced (+Dox) or uninduced (-Dox) for *Gata6* expression and cultured in 10 ng/ml Activin A with 0, 1, 3, 6, or 9 μM CHIR to modulate the WNT signaling pathway. Samples were analyzed at day 4.75. (B) Flow cytometry quantification for specification of mesoderm sub-type as in (A) except with culture in the presence of 75 ng/ml Activin A. (C) Volcano plot mapping changes in transcript levels at day 4 comparing RNA-seq of cells induced to express *Gata6* and exposed to DMSO or 6 μM CHIR at day 3. Note WNT-mediated down-regulation of DE markers and up-regulation of mesoderm markers. (D) Volcano plot mapping changes in transcript levels at day 4 comparing RNA-seq of cells exposed to 6 μM CHIR that were (+Dox) or were not (-Dox) induced to express *Gata6*. Note *Gata6*-mediated up-regulation of lateral plate mesoderm and down-regulation of paraxial and somitic mesoderm markers. (E) Comparison of the RNA-seq data from the samples described in (D) for the gene sets shown to be co-regulated by Activin and WNT signaling by Estaras et al., 2015. The pie chart shows that 90/117 (37 genes were not expressed or had less than 30 base mean counts) are significantly changed for expression levels by induction of *Gata6* and these patterns are illustrated by heat maps on the right. (F) Flow cytometry was used to quantify relative levels of non-DLL1 (top) or DLL1+ mesoderm sub-types at day 4 comparing WT or *Gata6*^-/-^ cells cultured in 75 ng/ml Activin A and exposed at day 3 to 0 (DMSO), 3 or 6 μM CHIR (CH3, CH6, respectively). (G) Gene expression analysis by RT-qPCR to measure relative transcript levels at day 4 of representative paraxial (top) or lateral (bottom) mesoderm markers for WT or *Gata6*^-/-^ cells cultured as in (F). Data in F and G are presented as mean ± SEM. Statistical significance in F and G was evaluated by unpaired, two-tailed Student’s t test for two-group comparisons

Comparison of the transcriptome of cells under +Dox/6 μM CHIR or +Dox/DMSO conditions confirms the WNT-mediated down-regulation of DE markers and the upregulation of ME markers (Figure 5C). Additionally, analysis of cells in +Dox/6 μM CHIR and -Dox/6μM CHIR reveals that Dox-induced expression of *Gata6*, when WNT is activated, upregulates markers associated with the lateral plate ME (Figure 5D). These markers include *Eomes*, *Mixl1*, and *FoxH1*, which are known to drive a highly conserved regulatory program for lateral plate ME (Prummel et al., 2019). Importantly, consistent with the flow cytometry data, *Gata6* down-regulates the expression levels of several drivers and markers of paraxial ME (*Tbx6*, *Msgn1*, *FoxC2*, *Lfng*, and *Aldh1A2*), which are otherwise enhanced by WNT activation (Figure 5D). GSEA analysis also reveals upregulation of multiple gene signatures associated with lateral plate ME and its derivatives following Dox-induced expression of *Gata6* and WNT stimulation. In contrast, several paraxial and somitic ME signatures are down-regulated (Figure S5E). Analysis of previously categorized WNT target gene sets demonstrates the significant regulatory role of *Gata6* in WNT-controlled networks, as evidenced by extensive transcriptional changes in genes either directly induced or repressed by WNT (Zhang et al., 2013) (Figure S5F and S5G). Furthermore, the expression analysis of 84 components of the WNT pathway, including ligands, receptors, effectors, positive and negative regulators, co-activators, and co-repressors, revealed that 54 of these components (64.3%) exhibited significant changes following *Gata6* expression (Figure S5H). These changes include the upregulation of *Fzd7* and the downregulation of *Fzd2*, genes shown to direct lateral or paraxial ME differentiation, respectively (Chidiac et al., 2025). Notably, while the induction of WNT direct targets by *Gata6* is associated with the induction of lateral plate ME but not DE (Figure S5I), the simultaneous induction of both DE and lateral plate ME by *Gata6* correlates with increased expression of *Nodal* and its direct targets (Figure S5J). Additionally, *Gata6* regulates the expression of genes co-regulated by Activin A and WNT (Estarás et al., 2015) (Figure 5E). Together, these results indicate that the role of *Gata6* in ME patterning arises from its simultaneous transcriptional regulation of WNT and NODAL signaling pathways.

The dual addition of CHIR and XAV has been shown to inhibit the nuclear transcriptional activity of β-Catenin while preserving its cytoplasmic functions (Kim et al., 2013). Using this combination, we found that ME specification, as well as transcription of *Axin2* and ME genes, are down-regulated in cells with or without Dox-induction of *Gata6* (Figure S6A-S6D), indicating that ME generation of cells expressing *Gata6* is also dependent on the transcriptional activity of β-Catenin. In addition, adding XAV with CHIR restored the DE population induced by *Gata6*, further confirming that *Gata6* renders DE progenitors independent of WNT signaling (Figure S6E). Taken together, these results indicate that expression of *Gata6* not only represses the WNT canonical pathway but also modifies its transcriptional output when WNT signaling activity is increased.

We showed that WNT signaling activity in the absence of *Gata6* persists at a high level during Activin-induced DE differentiation, as indicated by the expression levels of *Axin2* (Figure 4C). To examine the response to ME induction by WNT stimulation in the absence of *Gata6*, we analyzed the response of *Gata6^-/-^* cells to CHIR. As expected, based on our results with WT cells, increasing concentrations of CHIR resulted in a decrease in DE specification and an increase in ME progenitors (Figure S6F). In addition, the higher the CHIR concentration, the higher the yield of paraxial ME (Figure S6F). Notably, the type of ME progenitor generated by 3 μM or 6 μM CHIR differs between WT and *Gata6^-/-^* cells. While DLL1+ ME is significantly increased in *Gata6^-/-^* cells at 3 μM or 6 μM CHIR, non-DLL1 ME is significantly decreased, suggesting that WNT activation in the absence of *Gata6* biases fate toward paraxial ME (Figure 5F and S6G). Analysis of paraxial and lateral ME markers by qPCR confirmed the upregulation of markers of paraxial ME and down-regulation of markers associated with lateral plate ME in *Gata6^-/-^* cells after WNT activation (Figure 5G). These results show that *Gata6* regulates WNT signaling during ME specification to promote lateral versus paraxial ME fate.

### Gata6 Regulates *Eomes* and T/*Bra* Expression during both Endoderm and Mesoderm Specification

Although expression of *Eomes* and *T*/*Bra* overlaps initially in the primitive streak during mesendoderm formation, during lineage specification, *Eomes* antagonizes *T*/Bra to ensure the specification of DE and anterior ME (Schüle et al., 2023). Subsequently, the lack of expression of *Eomes* in more posterior regions allows *T/Bra* to drive axial, paraxial, and other posterior ME fates (Schüle et al., 2023). As *Gata4/6* expression induced a rapid increase in *Eomes* expression and decrease in *T*/*Bra* expression and modulates the gene networks regulated by both transcription factors (Figure 3D-G, S3D, and S3G), we hypothesized that *Gata6* regulates lineage segregation in cooperation with NODAL and WNT signaling through regulation of *Eomes* and *T*/*Bra*. The expression levels of *Eomes* and *T*/*Bra* were therefore evaluated by qPCR under conditions that induce DE (+Dox/DMSO), paraxial ME (-Dox/6 μM CHIR), or lateral ME (+Dox/6μM CHIR) (Figure 4G, 5A-B, and S5C-D). Increased expression levels of *Eomes* are associated with conditions that generate DE or lateral plate ME, while increased expression level of *T/Bra* is associated with both paraxial and lateral ME (Figure 6A and 6B). Notably, paraxial ME is associated with a faster and higher increase in *T/Bra* expression levels that subsided by 24 hr, at which point the expression level is similar as seen for lateral plate ME. In addition, we used flow cytometry to evaluate the effect of Dox-induced *Gata6* expression on BRACHYURY expression after WNT activation with CHIR at 3 μM or 6 μM. In cells without Dox, 3 μM or 6 μM CHIR generates paraxial ME (Figure 5A-B and S5C-D), and BRACHYURY expression is induced to similar levels (Figure S7A). In contrast, in +Dox conditions, BRACHYURY expression reaches similar levels to -Dox cells only in the presence of 6 μM CHIR (which in these conditions induces lateral plate-like ME) but not at 3 μM (Figure S7A), conditions in which *Gata6* instead promotes DE progenitors (Figure 4G, 5A-B, and S5C-D). This further confirms that *Gata6* inhibition of *T/Bra* is associated with the capacity to direct specification of DE.

**Figure 6.**
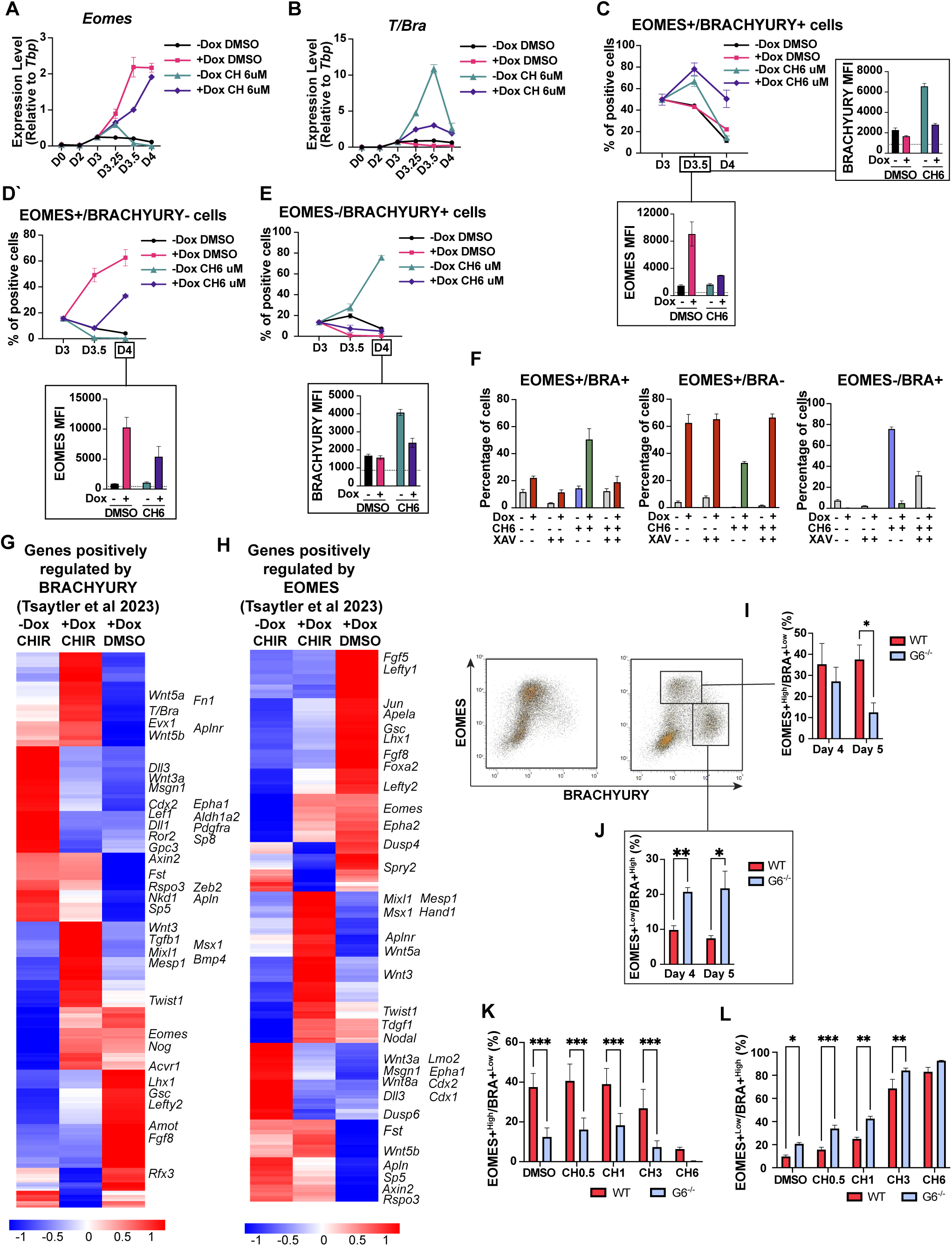
*Gata6* regulates both *Eomes* and *T*/*Bra* during endoderm and mesoderm specification. (A, B) Gene expression analysis by RT-qPCR at indicated time points to measure relative levels of *Eomes* (A) or *T/Bra* (B) in cells cultured in 10 ng/ml Activin A and induced by *Gata6* to specify DE (+Dox/DMSO), paraxial ME (-Dox/6 μM CHIR, CH), or lateral ME (+Dox/6 μM CHIR, CH), compared to those uninduced for *Gata6* or WNT (- Dox, DMSO). (C-E) Quantification by flow cytometry for the percentage of double-positive EOMES+/BRACHYURY+ cells (C), single-positive EOMES+ cells (D) and single-positive BRACHURY+ cells (E) at indicated time points following induction for *Gata6* expression at day 3 with Dox (or controls without Dox) and addition of either DMSO or 6 μM CHIR. MFI for each marker in positive populations is shown in associated graphs at day 3.5 (C) or day 4 (D, E). (F) Quantification by flow cytometry at day 4 as described for (C-E) except with or without addition of the WNT inhibitor XAV at day 3. Color of the bar represents predominant fate generated by each condition: endoderm (red), lateral mesoderm (green) and paraxial mesoderm (purple). White represents undefined lineage. (G, H) Heat maps based on RNA-seq at day 4 following addition of CHIR to activate WNT signaling in the absence (-Dox) or presence (+Dox) of Dox to induce expression of *Gata6* compared to Dox-induction of *Gata6* alone (+Dox, DMSO) evaluating the expression of genes positively regulated by BRACHYURY (G) and EOMES (H) as defined previously (Tsaytler et al., 2023) (I, J) Flow cytometry to quantify EOMES+ and BRACHYURY+ cells at day 4 or day 5 in WT or *Gata6*^-/-^ cells induced for DE specification with 75 ng/ml Activin A. Quantification is provided for EOMES+^High^/BRA+^Low^ (I) or EOMES+^Low^/BRA+^High^ (J). (K, L) Flow cytometry as in (I, J) except including 0 (DMSO) or 0.5, 1, 3, or 6 μM CHIR (CH). Cells were analyzed at day 4 (L) and day 5 (K). *n* ≥ 3 biologically independent experiments. Data in A-F and I-L are presented as mean ± SEM. Statistical significance in I and J was evaluated by unpaired, two-tailed Student’s t test for two-group comparisons. Statistical significance in B-F and K-L was evaluated with two-way ANOVA followed by Tukey’s test for multiple comparisons and significance for all these tests and comparisons is found in Table S3.

We also evaluated the co-expression of BRACHYURY and EOMES by flow cytometry. At day 3, before Dox-induced *Gata6* expression or CHIR addition, approximately 50% of cells co-express BRACHYURY and EOMES. Without Dox or CHIR, these progenitors transition to a double negative population by Day 4 (Figure 6C and S7B). 12 hr after Dox-induced *Gata6* expression (without CHIR), the percentage of EOMES^+^/BRACHYURY^+^ (E+/B+) double-positive cells is slightly reduced, and they display high levels of EOMES and low levels of BRACHYURY, as indicated by Median Fluorescence Intensity (MFI) (Figure 6C and S7B). 12 hr later (24 hr after Dox addition), most progenitors have transitioned to an EOMES^+^/BRACHYURY^-^ (E+/B-) single positive population expressing even higher EOMES levels (Figure 6D and S7B). In contrast, in 6 μM CHIR conditions, there is an initial increase in the number of E+/B+ cells at 12 hr regardless of Dox that is maintained up to 24 hr only in cells induced with Dox to express *Gata6* (Figure 6C). Of note, at 12 hr, the E+/B+ progenitors in -Dox/6 μM CHIR conditions have higher levels of BRACHYURY, while the same progenitors in +Dox/6 μM CHIR have lower levels (Figure 6C and S7B). By 24 hr, most progenitors in -Dox/6 μM CHIR conditions are EOMES^-^/BRACHYURY^+^ (E-B+) single-positive cells expressing the highest BRACHYURY levels for any conditions tested (Figure 6E). In contrast, at 24 hr in +Dox/6 μM CHIR conditions, around 80% of the population expresses EOMES, with half also expressing BRACHYURY (Figure 6C and 6D). One important distinction is that at 24 hr, the E+/B-cells in +Dox/6 μM CHIR expressed lower levels of EOMES compared to E+/B-cells in +Dox/DMSO conditions (Figure 6D and S7B).

Strikingly, when a WNT inhibitor is added, the E+/B+ population induced by +Dox/6 μM CHIR was replaced by an E+/B-population that expresses higher levels of EOMES (Figure 6F and S7C) and correlates with restoration of DE fate and repression of lateral ME (Figure S6C-S6E). Similarly, WNT inhibition in the -Dox/CHIR 6 μM conditions results in a significant decrease in the number of E-/B+ cells and reduced BRACHYURY MFI (Figure 6F and S7D), which correlates with the loss of paraxial ME specification observed (Figure S6B-S6D). Next, we compared the expression of positively regulated targets of EOMES and BRACHYURY in different ME types that were previously described (Faial et al., 2015; Harland et al., 2021; Koch et al., 2017; Tsaytler et al., 2023). WNT activation alone enhances the expression levels of multiple well-known drivers and markers of paraxial ME (*Wnt3a Msgn1*, *Cdx2*, *Alh1A2*, *Dll1*). In contrast, WNT activation along with Dox-induced expression of *Gata6* represses the expression of these targets and activates instead the expression of the lateral plate ME markers *Bmp4*, *Hand1*, *Tbx3*, *FoxF1*, and *Mesp1* (Figure 6G, 6H and S7E-S7G). These data indicate that *Gata6* promotes *Eomes* expression and represses *T/Bra* expression to specify DE. In contrast, WNT activation plays an opposing role, promoting *T/Bra* expression and repressing *Eomes* to drive the generation of paraxial ME. Furthermore, simultaneous activation of *Gata6* expression with WNT activation promotes *Eomes* and *T/Bra* co-expression to promote lateral-like ME fate.

Finally, to explore if this represents a normal function of *Gata6*, we compared the dynamics of EOMES and BRACHYURY expression from Day 2 to Day 5 during DE specification with a high dose of Activin A in the presence or absence of endogenous GATA6. In WT cells, EOMES expression initiates around Day 3 and increases to generate a population with high levels of EOMES and low levels of BRACHYURY (E+^High^/B+^Low)^ maintained up to Day 5 (Figure 6I and S8A). In contrast, although the number of E+^High^/B+^Low^ progenitors is not significantly different at day 4, a significantly lower number of E+^High^/B+^Low^ progenitors are derived by *Gata6^-/-^* cells at Day 5 (Figure 6I ad S8A), indicating a failure to maintain the expression of *Eomes* in the absence of GATA6 during the transition to DE fate. Furthermore, a distinct population expressing high levels of BRACHYURY and low levels of EOMES (E+^Low^/ B+^High^) was increased in the absence of GATA6 (Figure 6J and S8A). While the percentage of E+^High^/B+^Low^ cells on day 5 corresponds roughly to the same number of DE progenitors evaluated simultaneously in the experiment (DMSO in Figure S8E), the number of E+^Low^/ B+^High^ cells on Day 4 or Day 5 does not correlate to the number of ME progenitors (Figure S8B). Despite this, *Gata6^-/-^* cells display an increase in the expression of multiple ME markers, most of which are characteristic of paraxial ME (Figure S8C). Although the level of expression of ME markers is low when compared to conditions that directly specify ME (Figure 5G), the observation suggests that misregulation of *T/Bra* in the absence of GATA6 may underlie the differentiation bias of *Gata6^-/-^* cells toward paraxial ME fate in response to WNT activation (Figure 5F, 5G and S6F-G).

To test this, we evaluated the levels of EOMES and BRACHYURY in *Gata6^-/-^* cells in response to WNT activation with CHIR. Increasing concentrations of CHIR are associated with a decrease in the E+^High^/B+^Low^ population with a concomitant increase in the E+^Low^/ B+^High^ population (Figure S8D). The number of E+^High^/B+^Low^ cells peaks at Day 5 and correlates with the number of DE progenitors that decrease as the CHIR concentration increases (Figure S8E). In contrast, the E+^Low^/ B+^High^ population, which peaks at Day 4, correlates with the amount of ME generated (Figure S8F). Compared to WT, *Gata6^-/-^* cells generate a significantly lower percentage of E+^High^/B+^Low^ cells at all CHIR concentrations (Figure 6K and S8E). On the other hand, CHIR induces a higher percentage of E+^Low^/ B+^High^ cells at Day 4 in *Gata6^-/-^* cells correlating with the increased percentage of paraxial ME cells (Figure 5F, 6L, and S8F), demonstrating that the absence of GATA6 enhances the expression of *T/Bra* in response to WNT activation. Altogether, our results indicate that during DE specification, GATA6 is necessary for the upregulation and maintenance of *Eomes* and the downregulation of *T/Bra*. Furthermore, defective regulation of *T/Bra* in the absence of GATA6 is associated with a loss of lateral mesoderm capacity and a differentiation bias toward paraxial ME specification.

## DISCUSSION

The wide cellular diversity of a developing triplobastic organism arises from the three primary germ layers specified at the onset of gastrulation when the NODAL, WNT, and BMP signaling pathways converge to induce the PS (Tam and Loebel, 2007). During PS induction, these signaling pathways define a transient mesendoderm stage that will segregate to form the DE and ME lineages. How these mesendoderm progenitors developing in close spatiotemporal proximity interpret the same signaling cues to segregate into multiple fates remains a fundamental but unresolved question in early development. Using *Gata6*-inducible mouse embryonic stem cells, we demonstrate that *Gata6* expression during mesendoderm exit drives the DE GRN while simultaneously repressing the ME gene network regulated by WNT signaling. Unexpectedly, we found that upon activation of the WNT signaling pathway, GATA6 drives transcriptional changes in the expression of multiple components of the WNT pathway and a switch in the transcriptional output to promote lateral plate ME fate at the expense of paraxial ME fate. The different fate outcomes are correlated with the regulation of *Eomes* and *T/Bra* by GATA6 and WNT such that in DE, GATA6 upregulates EOMES and downregulates *T* expression and WNT signaling; in lateral plate ME, Gata6 upregulates *Eomes* and WNT upregulates *T*; and in posterior ME high WNT activity over-rides GATA6 and NODAL. Furthermore, we found that the absence of GATA6 results in a failure to deploy the DE GRN and produces a bias in ME differentiation towards paraxial ME. Thus, we define a new role for *Gata6* in the dual regulation of the transition from mesendoderm to both DE and ME, which helps explain the pleiotropic impact of patients with *GATA6* mutations (Škorić-Milosavljević et al., 2019; Tuhan et al., 2015; Yasuhara et al., 2024).

Our study suggests that loss of *Gata6* disrupts a positive feedback loop between EOMES and GATA6 that is integral to both DE and ME GRNs. Furthermore, our evidence indicates that GATA6 mediates some functions previously attributed to EOMES. Two distinct functions have been proposed for EOMES early in development. The initial activity of EOMES during exit from pluripotency and entry into the mesendoderm stage does not require transcriptional activity and involves de novo establishment of the accessible mesendoderm enhancer landscape (Schröder et al., 2023; Schüle et al., 2023). A subsequent lineage specification role for EOMES relies on transcriptional activity regulated by NODAL and WNT and is co-regulated by GATA6 (Chia et al., 2019; Schüle et al., 2023). Additionally, GATA6 and EOMES were found to interact and co-regulate genes required for precardiac ME patterning in the anterior lateral ME (Bisson et al., 2024). Using an Activin differentiation protocol similar to the one used in our study, it was shown that EOMES indirectly represses *T/Bra* to drive anterior fates (Schüle et al., 2023) through additional unidentified factors (Schüle et al., 2023). Our finding that *Gata6* is essential for maintaining *Eomes* expression and downregulating *T/Bra* to progress through the mesendoderm stage helps explain this proposed EOMES function. The role of GATA6 in *T/Bra* downregulation and ME repression during DE differentiation is also supported by recent reports that GATA6 promotes DE lineage commitment in part by restricting accessibility at enhancers linked to alternative lineages (Heslop et al., 2022).

It has been proposed that NODAL initially induces both DE and ME genes, while both lineages are segregated by mutually repressive gene networks (Zorn and Wells, 2009). Our findings about the role of *Gata6* in DE and ME differentiation enhance our understanding of how the NODAL and WNT signaling pathways interact to drive the networks that segregate lineage fate. During DE specification, *Gata6* reinforces NODAL signaling to maintain and enhance *Eomes* expression and create a positive feedback loop that contributes to the deployment of the full DE GRN. Additionally, GATA6 negatively regulates WNT to repress the ME GRN. Furthermore, *Gata6* expression marks the point of WNT inhibition independence in DE differentiation, which was proposed to occur when mesendoderm progenitors downregulate *T/Bra* before the overt expression of *Sox17* in DE progenitors (Pour et al., 2022). During mesendoderm exit toward ME fate, high WNT activity represses the expression of *Nodal* and its targets, including *Eomes* and *Gata6*. This repression in turn releases inhibition of *T/Bra* expression, as has been shown in vivo (Schüle et al., 2023), leading to establishment of the paraxial ME GRN (Mariani et al., 2021). Our inducible ESC model allowed us to show that lateral plate ME is favored by sustained activity of both NODAL/GATA6 and WNT, which maintains the expression of *Eomes* and *T/Bra*, respectively. We envision that EOMES cooperates with GATA6 and BRACHYURY to modify the transcriptional outputs of NODAL and WNT and deploy the lateral plate ME GRN, including *Mesp1*, *Hand1*, *Tbx3*, *Isl1*, and *FoxF1*. This mechanism is supported by the extensive literature documenting the cooperation of BRACHYURY and EOMES, GATA6 and EOMES and all these transcription factors with SMAD effectors of the NODAL signaling pathway (Bisson et al., 2024; Chia et al., 2019; Faial et al., 2015; Kumar et al., 2024; Liu et al., 2011; Prummel et al., 2019; Schüle et al., 2023; Slagle et al., 2011; Teo et al., 2011; Tosic et al., 2019; Tsaytler et al., 2023).

During subsequent organogenesis, regulation of the WNT pathway by *Gata6* has been described in heart and lung development (Afouda et al., 2008; Alexandrovich et al., 2006; Bisson et al., 2024; Lee and Evans, 2019; Novikov and Evans, 2013; Tian et al., 2010; Weidenfeld et al., 2002; Zhang et al., 2008). During lung development and airway regeneration after injury, loss of *Gata6* leads to an increase in canonical WNT signaling, impairing the balance between progenitor expansion and epithelial differentiation (Zhang et al., 2008). Additionally, regulation of WNT signaling by *Gata6* has emerged as an important factor promoting pancreatic and colon cancer, and *Gata6* is a critical contributor to WNT independence in a pathway involved in the switch from classical subtype to basal-like/squamous subtype of pancreatic cancer (Seino et al., 2018; Tsuji et al., 2014; Whissell et al., 2014; Zhong et al., 2011, 2022). Recapitulation of critical developmental pathways in the regulation of progenitor cell expansion is required for tissue regeneration and is altered in tumorigenesis; thus, our findings that *Gata6* is a regulator of fate through modulation of WNT signaling helps explain how *Gata6* may also regulate similar processes in regeneration and disease.

## EXPERIMENTAL PROCEDURES

Detailed methods can be found in the supplemental information and are briefly described below.

### Cell lines, culture, and differentiation

Mouse embryonic stem cell lines engineered to allow conditional expression of *Gata4* or *Gata6* were maintained as previously described (Turbendian et al., 2013). ESCs were dissociated and aggregated for differentiation as embryoid bodies (EBs) as previously described (Gouon-Evans et al., 2006). The Gata4-inducible mesendoderm reporter was generated by targeting the *Gata4* cDNA into the tet-regulated promoter locus of the AinV/GFP-Bry/CD4-Foxa2 ESC line as previously described (Cheng et al., 2008; Holtzinger et al., 2010). For liver differentiation and mid-hindgut differentiation, day 5 EBs were dissociated with Accutase and plated at a ratio of 1:2, and differentiation was performed as previously described (Gouon-Evans et al., 2006; Loh et al., 2014) with modifications as detailed in the supplemental information.

### RNA isolation, RT-qPCR analysis, and Bulk RNA sequencing

Detailed methods for processing and analysis methods, as well as primer sequences, can be found in supplemental information and Table S1.

### Flow cytometry

For extracellular markers, EBs were harvested, dissociated, and stained with antibodies before being stained with Sytox blue for viability. For intracellular antigens, cells were fixed with 2% PFA before staining with antibodies. Samples were analyzed on an Attune Nxt (Thermofisher). See supplemental information for details on the staining and antibodies used.

### Western blotting

Proteins were extracted using RIPA buffer supplemented with 1x protease and phosphatase inhibitors (Pierce), and benzonase (Millipore). Lysates were centrifuged at 15,000 rpm for 5 min at 4°C. Total protein concentration was measured using the bicinchoninic acid protein assay kit (Pierce 23225). See supplemental methods for more details.

### Quantification and statistical analysis

All data is presented as mean ± SEM of independent biological replicates with n indicated in each figure legend. GraphPad Prism software was used for statistical analysis with two-tailed Student’s *t*-test when comparing two groups and one-way or two-way ANOVA when comparing three or more groups, as indicated in figure legends. Statistically significant p values were indicated in figures as *, p < 0.05; **, p < 0.01; ***, p < 0.001; ****, p < 0.0001. Non-significant comparisons are not indicated in figures.

## RESOURCE AVAILABILITY

### Lead contact

Requests for further information and resources should be directed to and will be fulfilled by the lead contact, Todd Evans.

### Materials availability

All unique/stable reagents generated in this study are available from the lead contact.

### Data and code availability

The accession number for the RNA-seq data reported in this paper is deposited at the Gene Expression Omnibus (GEO) Database: GSEXXXXX.

## ACKNOWLEDGEMENTS

We thank Gordon M. Keller from the University Health Network in Toronto and Valerie Gouon-Evans from Boston University for sharing the AinV/GFP-Bry/CD4-Foxa2 cell line. Stephen A. Duncan from the Medical University of South Carolina graciously shared the *Gata4*^-/-^, *Gata6*^-/-^, and *Gata4*^-/-^;*Gata6*^-/-^ cell lines generated by his group. We want to thank all members of the Evans laboratory for their constructive feedback throughout this study. This work was supported by a grant from the NIH to TE (R35 HL135778).

## AUTHOR CONTRIBUTIONS

MG designed and performed the experiments, analyzed the data, and wrote the manuscript. RW and NdS performed experiments, KB helped analyze data. TE contributed with funding acquisition, experimental design, interpretation of the data, and wrote the manuscript.

## Supplemental Figures and Text

**Figure S1.**
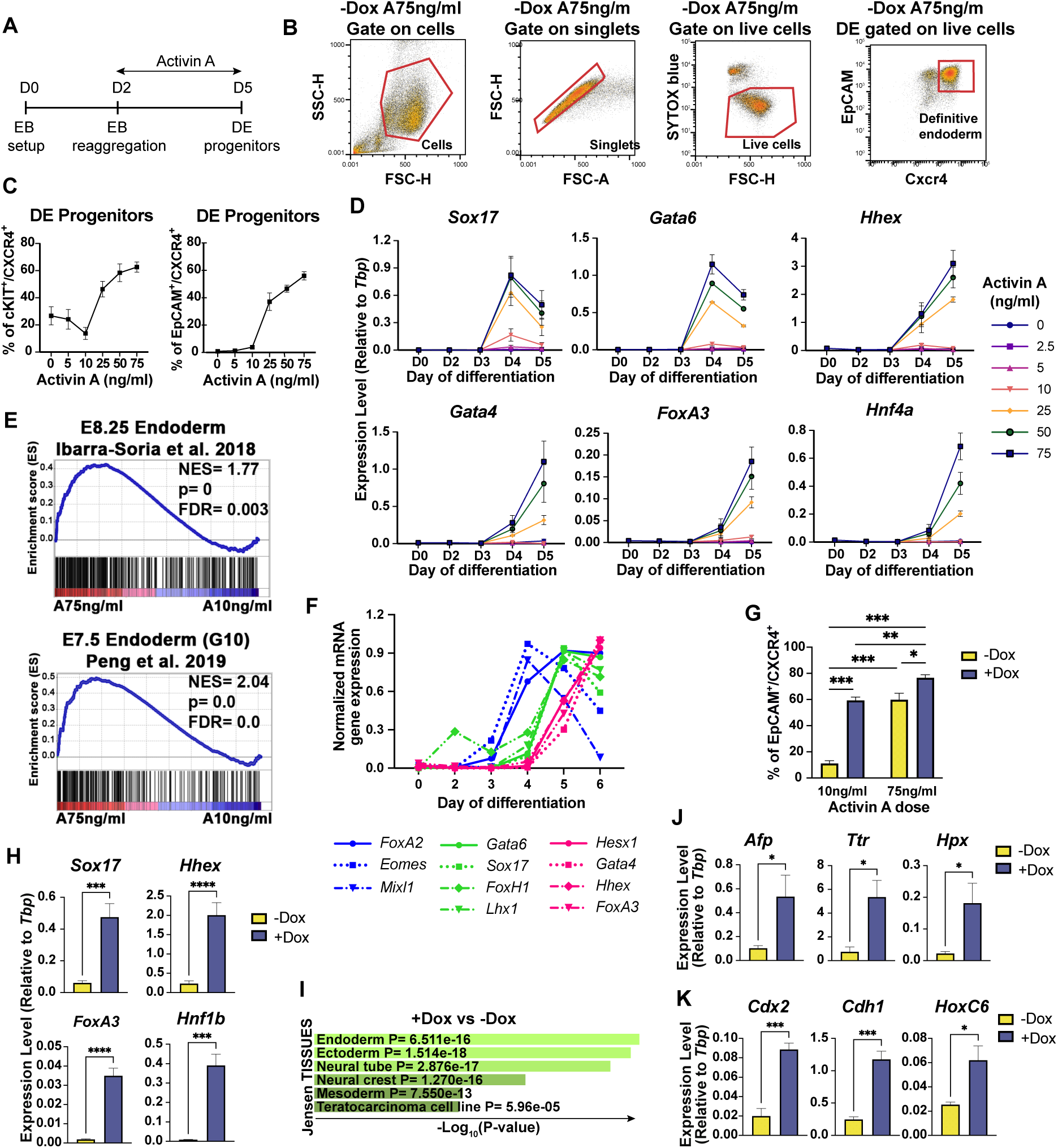
Expression of GATA4 or GATA6 can drive murine definitive endoderm specification, and GATA6 is required. Related to Figure 1. (A) Schematic diagram of the definitive endoderm (DE) directed differentiation protocol. Embryoid bodies (EBs) were set up on day 0 and reaggregated on day 2 in the presence of 75 ng/ml Activin A. (B) Representative gating strategy for identification of progenitors after differentiation. ESCs cultured without doxycycline (in experiments with inducible cell lines) or WT cells (in experiments with GATA KO cells) at day 5 of differentiation were used to define the gating strategy used throughout all DE flow cytometry experiments. To exclude debris, cells were gated from all events by analyzing FSC-H and SSC-H. Then, cells were analyzed by FSC-A and FSC-H scatter to exclude doublets. After gating on singlets, live cells for analysis of surface markers were gated as the Sytox Blue negative population. (C) Flow cytometry to quantify DE specification at day 5 comparing cKIT+/CXCR4+ or EpCAM+/CXCR4+ populations at different Activin A concentrations. (D) RT-qPCR analysis for key DE markers at the indicated day (D) of differentiation using various concentrations of Activin A showing activation at and above 25 ng/ml. (E) GSEA analysis of the transcriptome from day 4 RNA-seq data comparing 10 or 75 ng/ml Activin A shows enrichment with 75 ng/ml Activin A of DE signatures derived from single-cell RNA-seq of mouse embryos. (F) Temporal expression analysis by RT-qPCR of key TFs of the DE Gene Regulatory Network at the indicated day of differentiation using 75 ng/ml Activin A showing 3 waves of activation around day 3 (*FoxA2*, *Eomes*, *Mixl1*), day 4 (*Gata6*, *Sox17*, *FoxH1*, *Lhx1*), and day 5 (*Hesx1*, *Gata4*, *Hhex*, *FoxA3*). Data points represent the mean normalized to the maximum expression of each gene. (G) Flow cytometry analysis of DE progenitors at day 5 using the *Gata4*-inducible ESC line and culture conditions in either 10 ng/ml or 75 ng/ml Activin A and either DMSO or Dox added at day 3. (H) RT-qPCR analysis for key DE markers at day 5 using the *Gata4*-inducible ESC line cultured in 10 ng/ml Activin A and either DMSO or Dox added at day 3. (I) Gene list analysis of the *Gata6*-induced transcriptome using Enrichr shows a high enrichment of genes associated with DE development. (J, K) Following induced expression of *Gata4*, day 5 EBs were replated in hepatoblast (J) or mid-hindgut (K) differentiation conditions and evaluated by RT-qPCR analysis after 4 days (J) or 3 days (K) for expression of liver (J) or hindgut (K) markers. Statistical significance in H, J, and K was evaluated by unpaired, two-tailed Student’s t test for two-group comparisons. Statistical significance in G was evaluated with two-way ANOVA followed by Tukey’s test for multiple comparisons.

**Figure S2.**
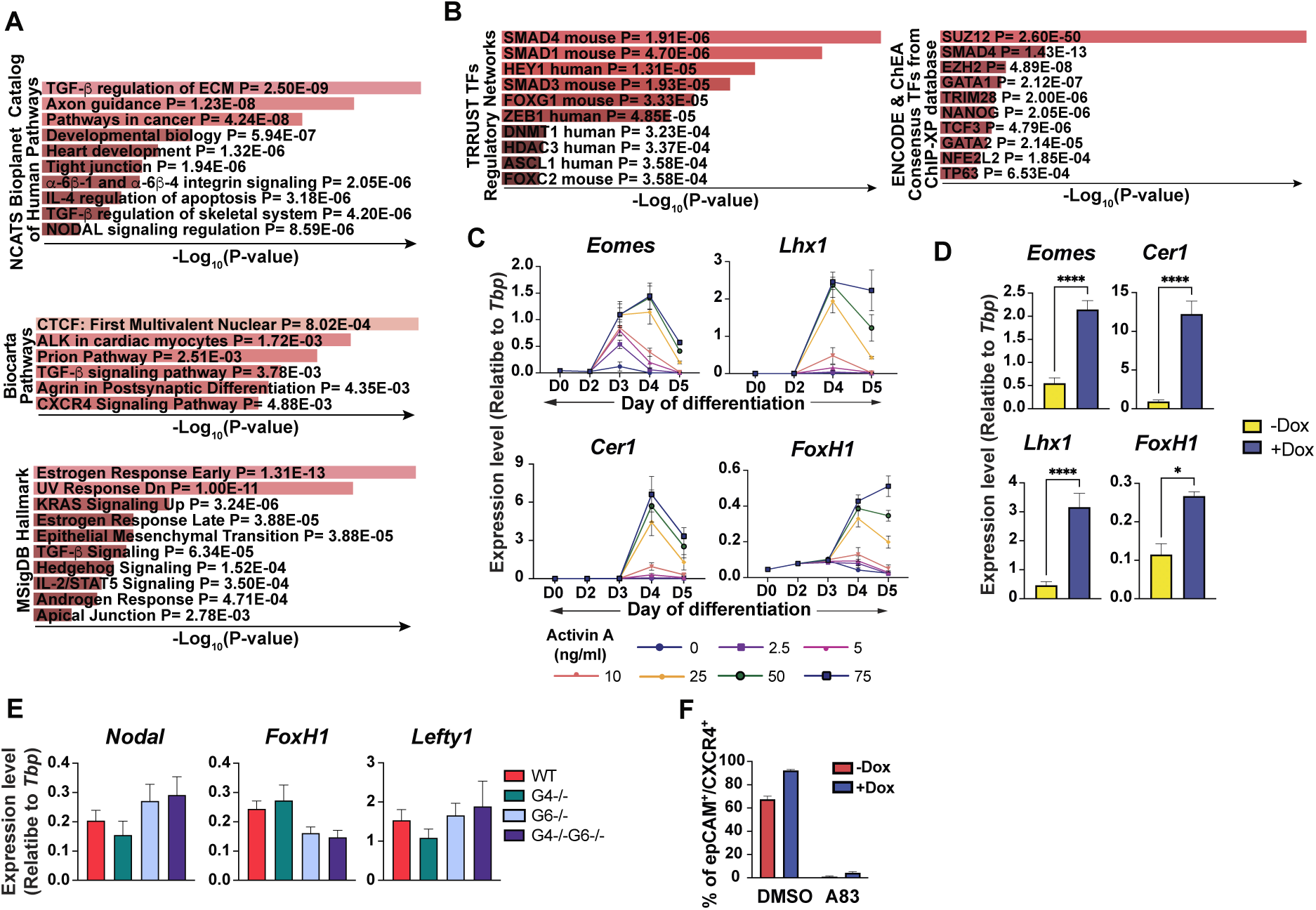
GATA6 reinforces the NODAL pathway. Related to Figure 2. (A, B) Enrichment analysis of the *Gata6*-induced transcriptome using RNA-seq data highlights gene sets associated with the TGFβ/NODAL signaling pathway (A) and SMAD2/3/4 (B). (C) RT-qPCR analysis for known direct SMAD2/3/4 target genes at the indicated day of differentiation using various concentrations (ng/ml) of Activin A. (D) RT-qPCR analysis for cells cultured in 10 ng/ml Activin A collected 24h after they were induced (+Dox) or not (-Dox) for expression of *Gata4*. Statistical significance was evaluated by unpaired, two-tailed Student’s t test for two-group comparisons. (E) RT-qPCR analysis at day 5 for key NODAL pathway regulators in WT or GATA mutant cells cultured in 75 ng/ml Activin A. Statistical significance was evaluated with two-way ANOVA followed by Tukey’s test for multiple comparisons. No statistically significant differences were found. (F) As quantified by flow cytometry at day 5, specification of DE by induction of *Gata6* expression (in 75 ng/ml Activin A) requires active Activin signaling, as it is blocked by addition of the A83-01 ALK4/5/7 type I receptor inhibitor.

**Figure S3.**
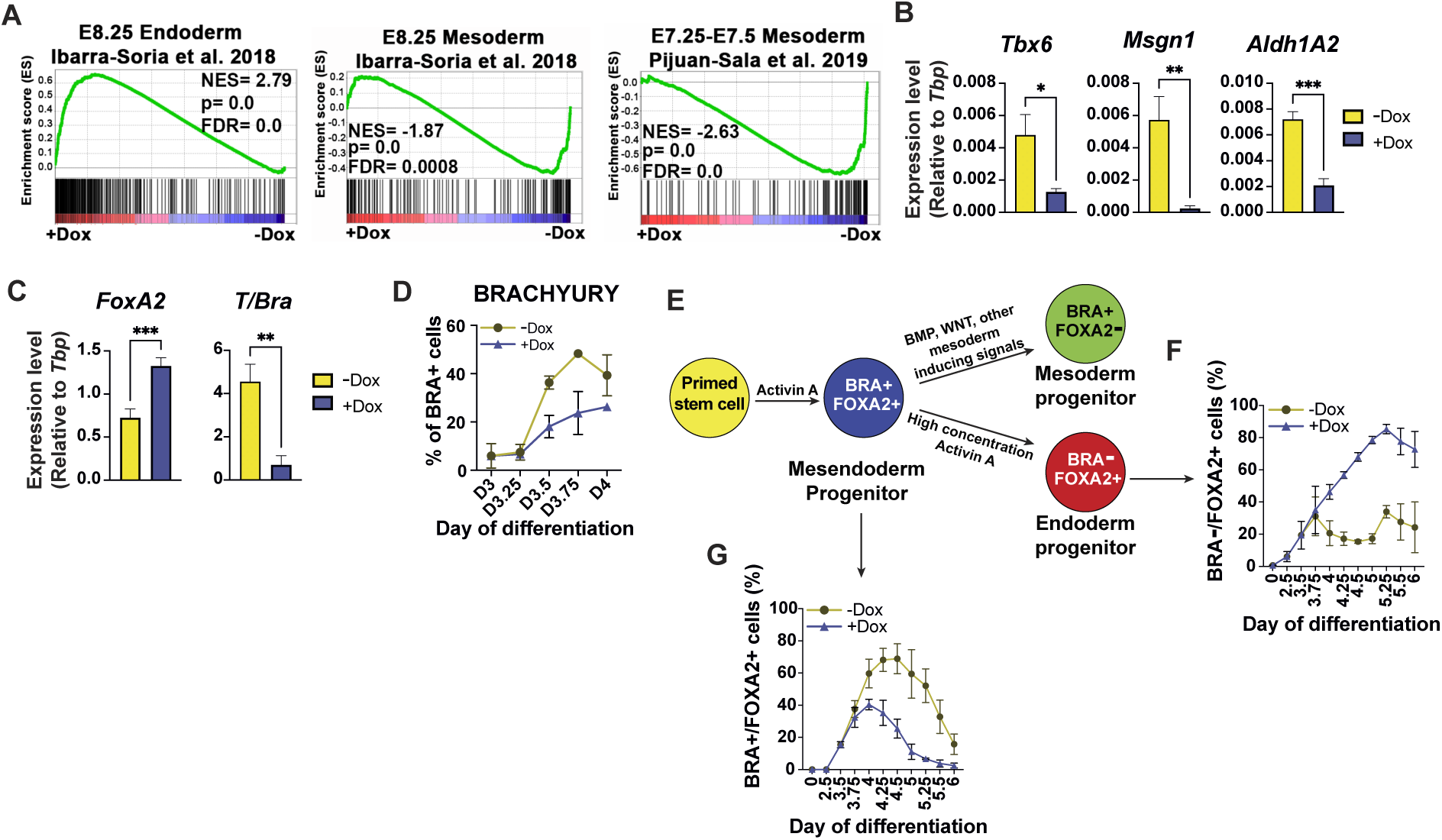
GATA factors drive mesendoderm cells to definitive endoderm fate. Related to Figure 3. (A) GSEA using RNA-seq data of cells at day 4 with or without Dox-induced expression of *Gata4* at low concentration of Activin A shows suppression of gene sets for mesoderm. (B) Gene expression analysis by RT-qPCR for key mesoderm drivers on day 4 of differentiation following Dox-induction of *Gata4* and culture in 10 ng/ml Activin A. (C) Gene expression analysis by RT-qPCR at day 4 of differentiation following Dox-induction of *Gata4* and culture in 10 ng/ml Activin A shows enhanced expression of *FoxA2* and down-regulation of *T/Bra*. (D) Flow cytometry to quantify the percentage of cells expressing BRACHURY at defined time points shows Dox-induction of *Gata4* expression reduces BRA+ cells. (E) Schematic illustrating of proposed lineage progression and expression of FOXA2 and BRACHYURY during the mesoderm and endoderm differentiation from mesendoderm progenitors (Gadue et al., 2006). (F, G) Flow cytometry results using the dual reporter line to follow BRA-GFP and FOXA2-CD4 expression when cultured in 10 ng/ml Activin A and induced (or not) with Dox at day 3 showing BRA-/FOXA2+ DE (F) and BRA+/FOXA2+ mesendoderm (G). Data are presented as mean ± SEM. Statistical significance in B and C was evaluated by unpaired, two-tailed Student’s t test for two-group comparisons.

**Figure S4.**
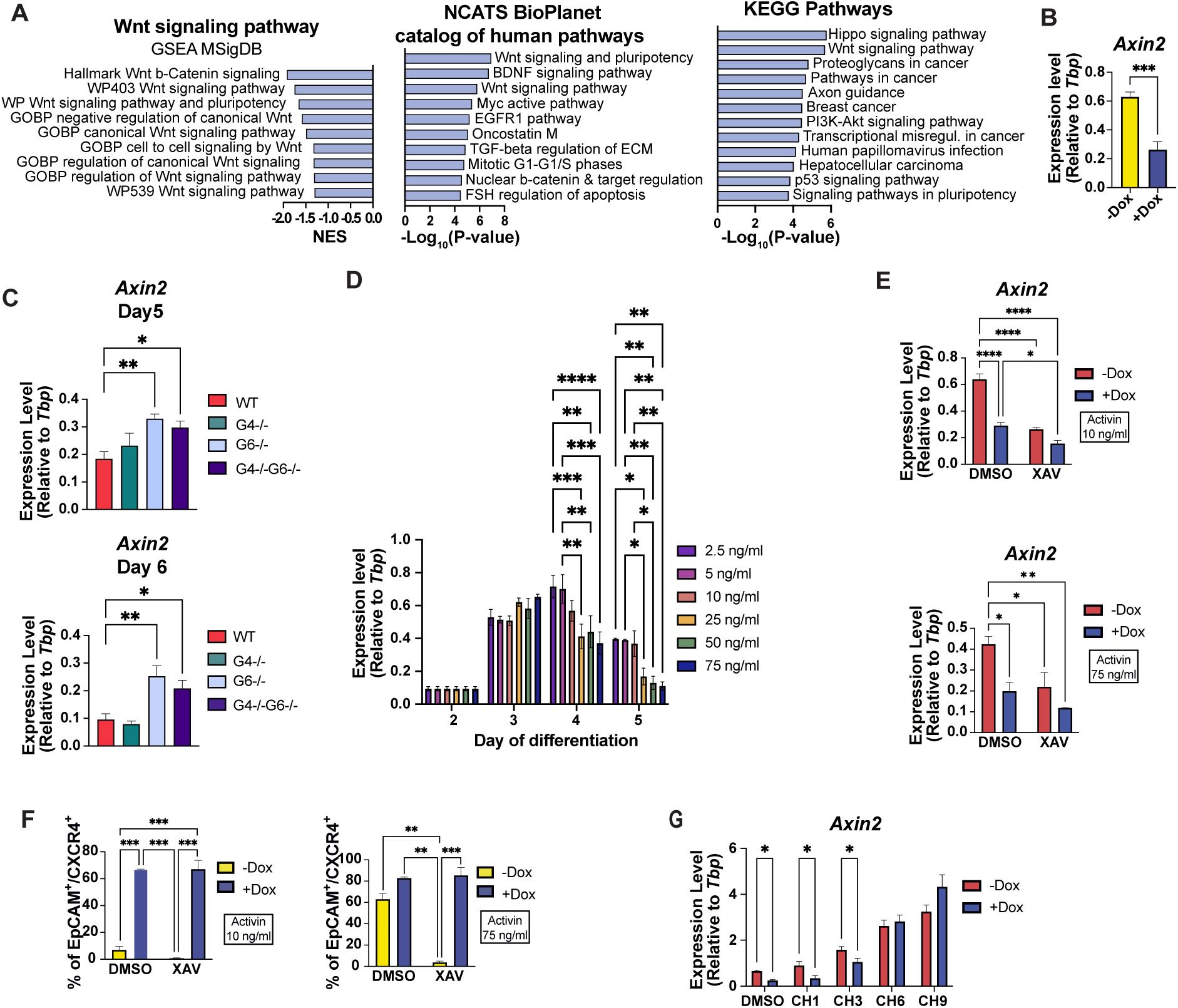
GATA6 negatively regulates WNT signaling and grants WNT independence during DE specification. Related to Figures 3 and 4. (A) GO terms and pathway enrichment analysis of differentially regulated genes at day 4 following Dox-induced *Gata4* expression shows WNT/β-catenin signaling as one of the main down-regulated pathways. (B) Gene expression analysis by RT-qPCR shows relative down-regulation of the canonical WNT early target gene *Axin2* at day 4 in cells induced to express *Gata4* during differentiation using 10 ng/ml Activin A. (C) Gene expression analysis by RT-qPCR to measure relative *Axin2* transcript levels at indicated time points in cells lacking GATA4, GATA6, or both proteins compared to WT cells during differentiation using 75 ng/ml Activin A. Only GATA6 is required to repress Axin transcript levels. Related to Figure 3C. (D) Gene expression analysis by RT-qPCR at indicated time points to measure relative *Axin2* transcript levels in WT cells induced by different concentrations of Activin A. Related to Figure 4D. (E) RT-qPCR analysis of *Axin2* transcript levels confirms the inhibitory effect of XAV in both - Dox and +Dox conditions and at 10 or 75 ng/ml Activin A. (F) Flow cytometry was used to measure DE specification at day 5 showing that induction of *Gata4* expression is sufficient to rescue DE fate from XAV-mediated WNT inhibition. (G) RT-qPCR analysis of *Axin2* transcript levels indicates that cells induced to express *Gata6* respond to the activation of the pathway but require a higher concentration of CHIR to achieve the same level of *Axin2* expression compared to cells not induced by Dox. Data in B-G are presented as mean ± SEM. *n* ≥ 3 biologically independent experiments. Statistical significance in B and G was evaluated by unpaired, two-tailed Student’s t test for two-group comparisons. Statistical significance in C was evaluated with one-way ANOVA followed by Dunnett’s test for multiple comparisons relative to WT. Statistical significance in D-F was evaluated with two-way ANOVA followed by Tukey’s test for multiple comparisons.

**Figure S5.**
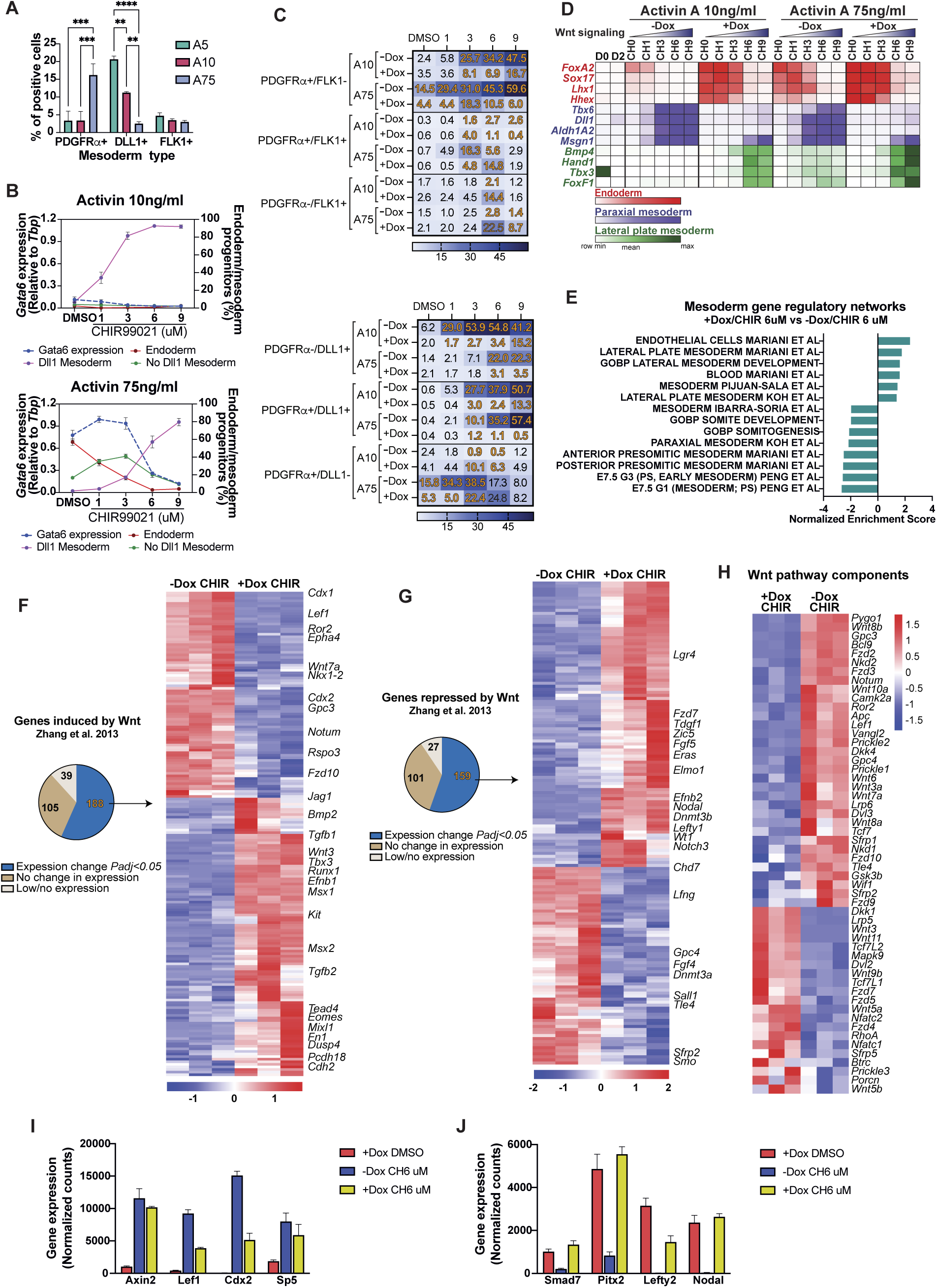
GATA6 cooperates with WNT signaling to specify mesoderm subtype. Related to Figure 5. (A) Quantification of flow cytometry at day 4.75 to measure the percentage of mesoderm sub-types based on PDGFRα, DLL1, and FLK1 expression, as impacted by the concentration of Activin A (5, 10, or 75 ng/ml). (B) WT cells were cultured in 10 ng/ml (top) or 75 ng/ml (bottom) Activin A and various concentrations of WNT activator CHIR99021, as indicated. Graphs chart the relative level of *Gata6* expression (based on RT-qPCR) correlating with relative percent of DE, DLL1+ mesoderm, or non-DLL mesoderm (based on flow cytometry), showing that *Gata6* levels are associated with Activin/CHIR that generates either a high yield of DE or non-DLL1 progenitors. Flow cytometry samples were collected at day 4.75 and RNA samples were collected at day 4. (C) Analysis of flow cytometry data at day 4.75 as in (B) for mesoderm sub-types (PDGFRα and FLK1, top; PDGFRα and DLL1, bottom) when cells were cultured with low (10 ng/ml) or high (75 ng/ml) Activin A plus increasing concentrations of CHIR and either treated at day 3 with DMSO (-Dox) or with Dox to induce expression of *Gata6*. With induction of *Gata6*, DLL1+ mesoderm is largely abrogated, but at high concentration of Activin A promotes generation of FLK1+ progenitors. (D) Analysis as in (B) except that cells were harvested at day 4 for RT-qPCR to evaluate markers for DE, paraxial mesoderm, and lateral plate mesoderm, for which relative expression levels are shown on the heat maps. Expression of *Gata6* promotes DE but with high concentrations of WNT (activated by CHIR at 6 or 9 μM) promotes lateral plate mesoderm at the expense of paraxial mesoderm. (E) GSEA based on RNA-seq at day 4 for cells cultured in 6 μM CHIR comparing those uninduced or induced (+Dox) to express *Gata6*, showing promotion of lateral plate mesoderm and repression of paraxial or somitic mesoderm signatures. (F, G) Heat maps show RNA-seq profiles from samples as in (E) for gene sets that are induced (F) or repressed (G) by WNT, showing that *Gata6* widely impacts WNT-controlled gene networks. (H) Heat map shows RNA-seq profiles for components of the WNT signaling pathway for which 54/84 are altered significantly by expression of *Gata6*. (I) Gene expression evaluated as normalized counts from RNA-seq at day 4 for known direct targets of WNT signaling. (J) Gene expression evaluated as normalized counts from RNA-seq at day 4 for known direct targets of NODAL signaling.

**Figure S6.**
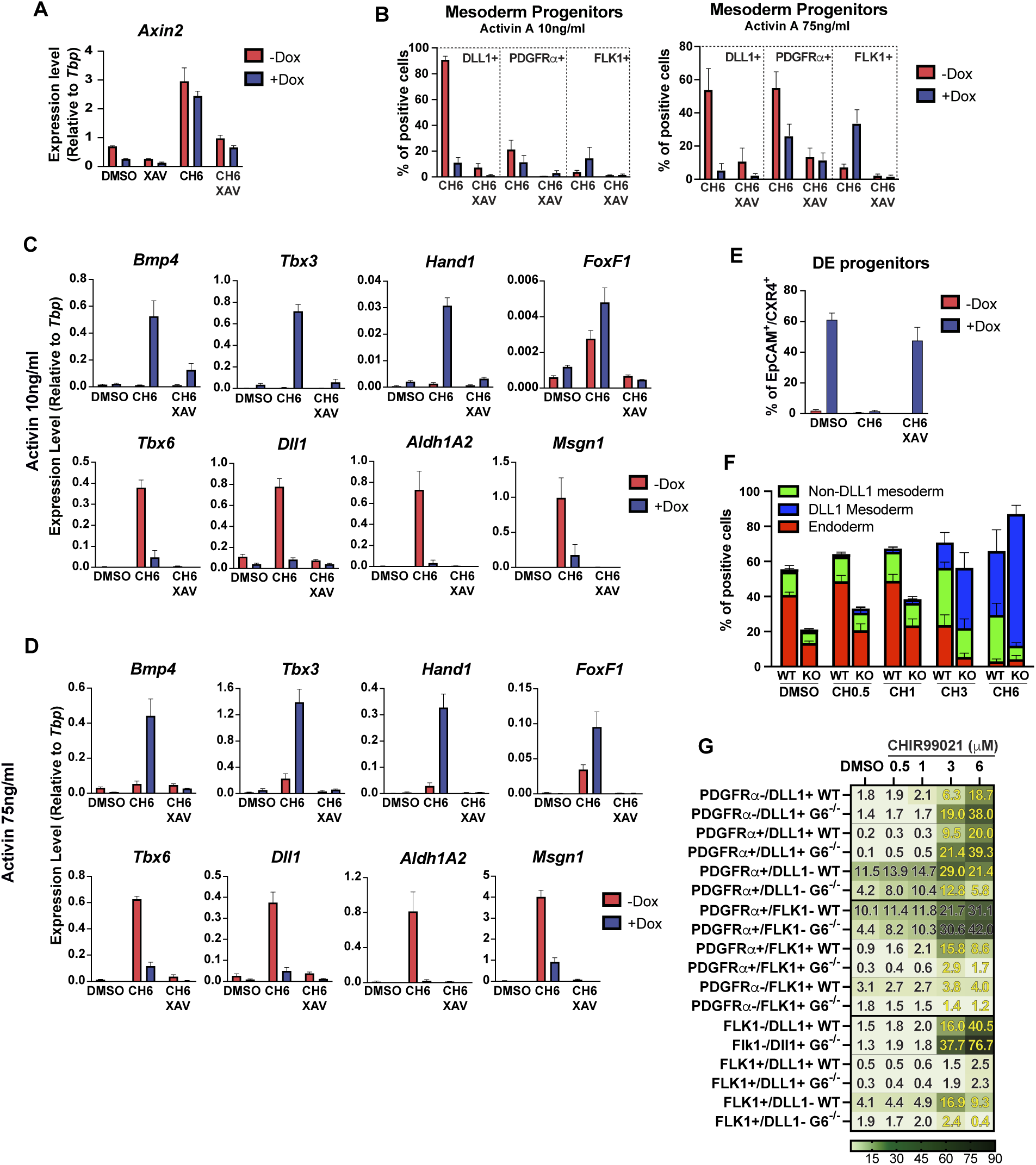
GATA6 cooperates with WNT signaling to specify mesoderm subtype. Related to Figure 5. (A-D) RT-qqPCR assays (A, C, D) or flow cytometry (B) were performed after differentiation in DMSO, XAV, 6 μM CHIR, or both XAV and CHIR, either without (-Dox) or with Dox-induction (+Dox) of *Gata6*. Assays quantified levels of WNT target *Axin2* (A), mesoderm sub-types (B) and mesoderm sub-type markers in 10 ng/l Activin A (C) or 75 ng/ml Activin A (D). (E) Flow cytometry to measure DE specification at day 5 showing that expression of *Gata6* (+Dox) promotes DE fate when XAV is added in cultures with 6 μM CHIR. (F) Flow cytometry was used to measure at day 5 the specification of DE (red) and DLL1+ (blue) or non-DLL mesoderm subtypes using WT or *Gata6*^-/-^ (KO) cells treated with 0, 0.5, 1, 3, or 6 μM CHIR as indicated. In cells lacking GATA6, CHIR promotes specification of DLL1+ mesoderm. (G) Flow cytometry data to quantify mesoderm sub-types comparing WT and *Gata6*^-/-^ cells cultured in various concentrations of CHIR, as indicated.

**Figure S7.**
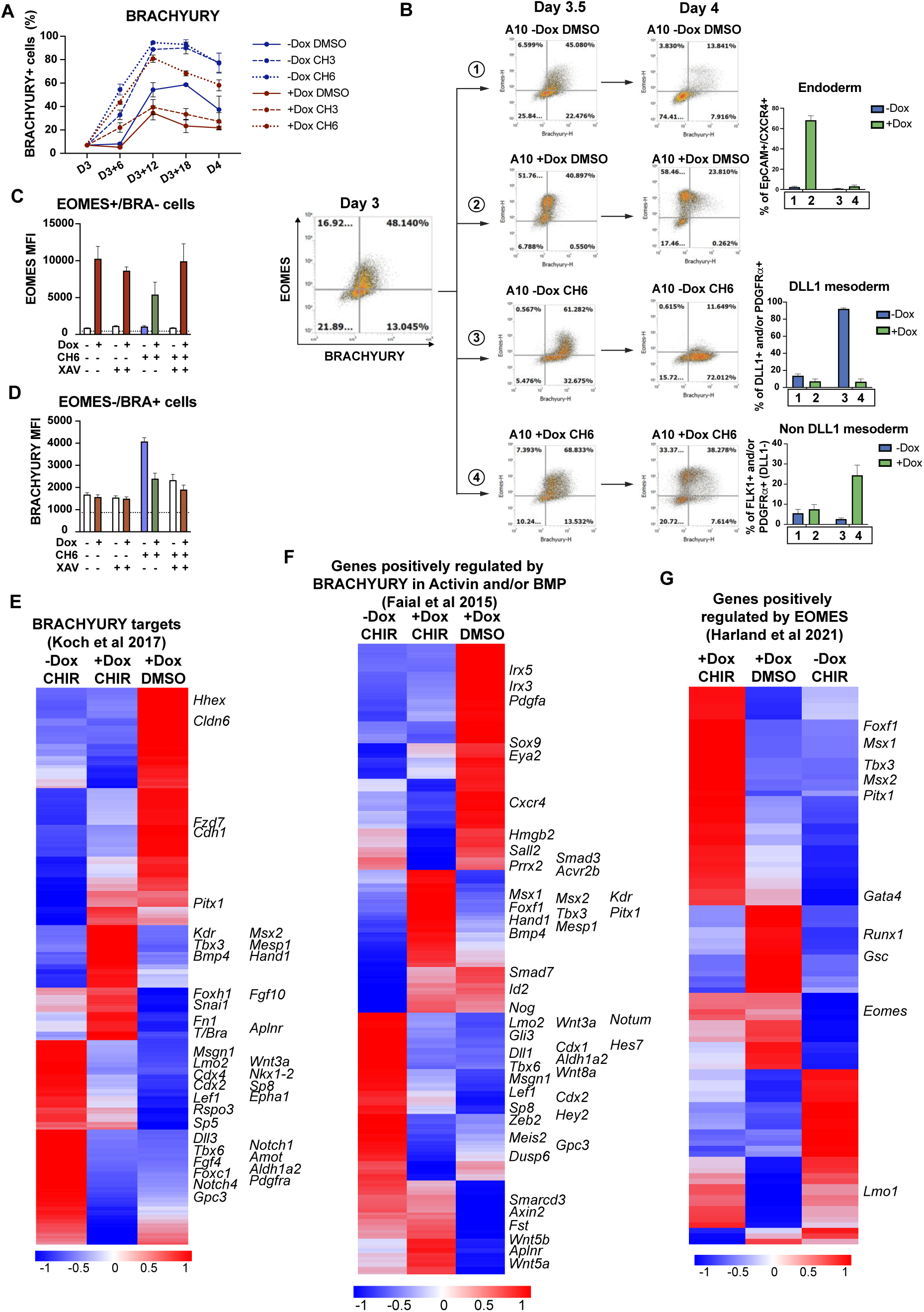
*Gata6* regulates *Eomes* and *T*/*Bra* during both endoderm and mesoderm specification. Related to Figure 6. (A) Flow cytometry was used to quantify the percentage of BRA+ cells every 6 hours from day 3 to day 4 for cells cultured in 10 ng/ml Activin A and 0, 3, or 6 μM CHIR and either uninduced (-Dox) or induced with Dox (+Dox) at day 3 to express *Gata6*. (B) Flow cytometry was used to quantify percentage of cells expressing EOMES and/or BRACHYURY initially at day 3 (left panel) and then at day 3.5 or 4 middle panels) following culture in 10 ng/ml Activin A, without or with 6 μM CHIR and without (-Dox) or with induction (+Dox) to express *Gata6*. Graphs on the far-right associate each of the four conditions with the generation of DE, DLL1+ mesoderm, and non-DLL1 mesoderm. Expression of *Gata6* alone drives DE (EOMES up, BRA down), CHIR alone drives DLL1+ mesoderm (EOMES down, BRA up), and *Gata6* expression in the presence of CHIR drives predominantly non-DLL mesoderm (EOMES and BRA both sustained). (C, D) MFI data for EOMES in EOMES+/BRA-cells (C) or BRACHYURY in EOMES-/BRA+ cells (D) quantified in Figure 6F. Color of the bar represents predominant fate generated by each condition: endoderm (red), lateral mesoderm (green) and paraxial mesoderm (purple). White represents undefined lineage. (E-G) Heat maps show transcriptomic profiles from RNA-seq data at day 4 for cells cultured in 6 μM CHIR (or DMSO) and induced with Dox at day 3 to express *Gata6* (or without Dox) to compare impact on genes regulated by BRACHYURY as defined by (E) Koch et al. (2017) and (F) Faial et al. (2015) or EOMES targets as defined by (G) Harland et al. 2021.

**Figure S8.**
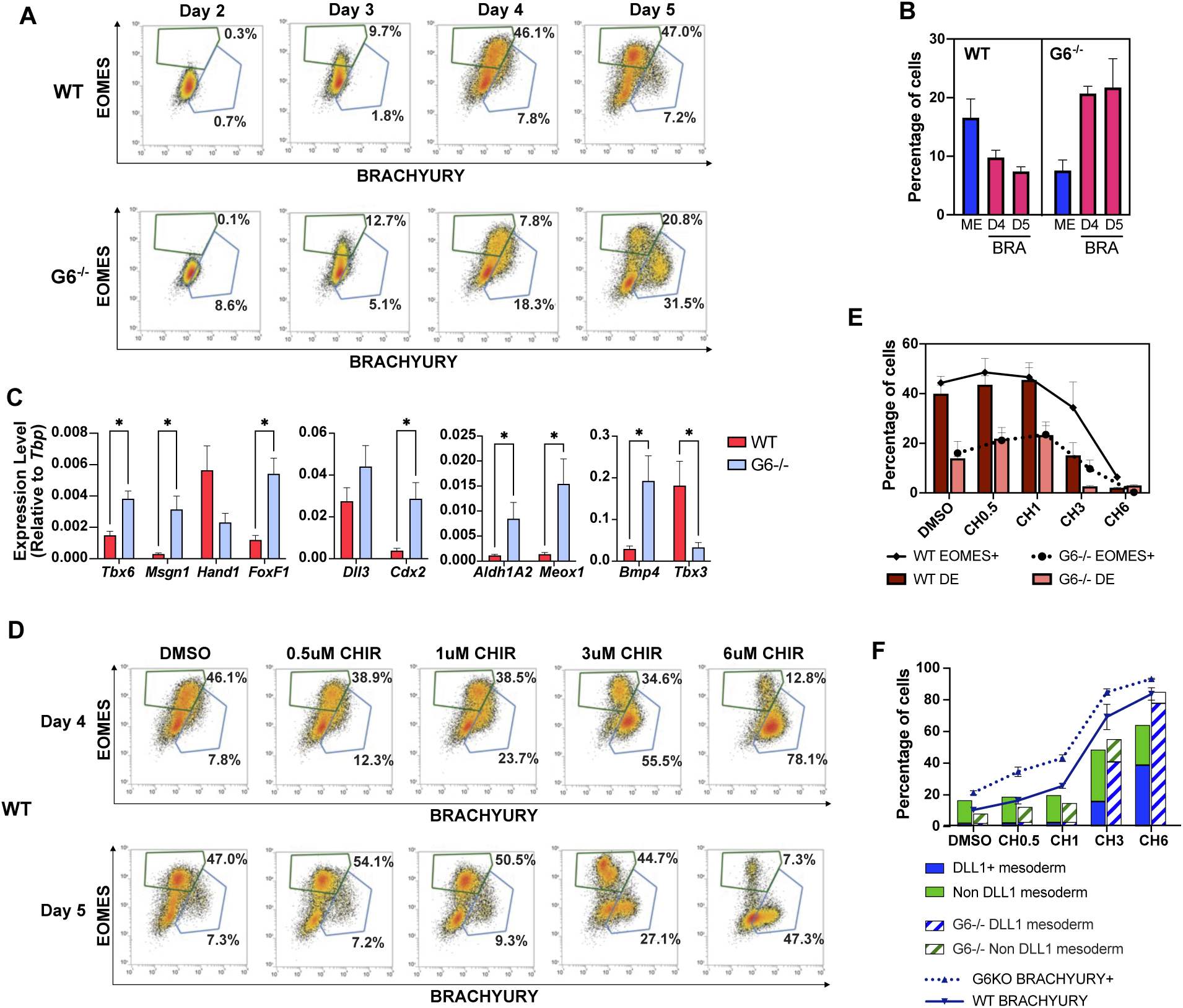
*Gata6* regulates *Eomes* and *T*/*Bra* during both endoderm and mesoderm specification. Related to Figure 6. (A) Flow cytometry was used to quantify EOMES and BRACHURY expressing cells at day 2-5 as indicated for WT or Gata6^-/-^ cells cultured in 75 ng/ml Activin A. Shown are representative flow plots. (B) BRA+ cells from (A) at day 4 or day 5 are shown relative to the percentage of DLL1+ and non-DLL mesoderm progenitors from the same samples at day 5 (ME). (C) RT-qPCR assays were used to quantify levels of ME markers in day 5 samples from (A) showing primarily increased levels of paraxial markers in the *Gata6*^-/-^ cells. (D) Flow cytometry was used to quantify in WT cells the percentage of EOMES and BRACHURY expressing cells at day 4 following culture in 75 ng/ml Activin A (as in A) except with the addition at day 3 of 0 (DMSO), 0.5, 1, 3, or 6 μM CHIR. Higher levels of CHIR substantially shift the progenitor population to BRA+ and EOMES-fate. (E) Flow cytometry data demonstrates a correlation between loss of EOMES+ cells and DE as cells are exposed in 75 ng/ml Activin A to increasing levels of CHIR, which is much enhanced in the absence of GATA6. (F) Flow cytometry data for mesoderm sub-types in WT or *Gata6*^-/-^ cells in 75 ng/ml Activin A following addition of 0 (DMSO), 0.5, 1, 3, or 6 μM CHIR. CHIR drives specification to ME fate and in the absence of GATA6 this is almost entirely DLL1+ paraxial mesoderm that is also BRA+.

## SUPPLEMENTAL METHODS

### Cell lines, culture, and differentiation

All cells were cultured under humidified normoxic conditions at 37°C and 5% CO_2_ in an air-jacketed incubator. Mouse embryonic stem cell lines engineered to allow conditional expression of Gata4 and Gata6 and the Gata4^-/-^, Gata6^-/-^, and Gata4 ^-/-^;Gata6^-/-^ cell lines have been previously described (Holtzinger et al., 2010; Molkentin et al., 1997; Turbendian et al., 2013; Zhao et al., 2008, 2005). The Gata4 inducible GFP-Bry/CD4-FoxA2 ES cell line was generated by targeting Gata4 cDNA into the tet-regulated promoter near the HPRT locus of the AinV/GFP-Bry/CD4-Foxa2 cells as previously described (Cheng et al., 2008; Holtzinger et al., 2010). Stably transfected cells were selected using increasing concentrations of G418 (Cellgro; up to 300 μg/ml). Individual clonal lines were selected by picking colonies. Proper integration of the transgene was confirmed by PCR analysis of genomic DNA using the following primers: fwd: 5′-CTAGATCTCGAAGGATCTGGAG-3′; rev: 5′-ATACTTTCTCGGCAGGAGCA-3′.

All cell lines were maintained on plates coated with 0.2% gelatin in 2i+LIF media composed of 25% DMEM (Corning 15-018-CV), 25% Ham’s F12 (Corning 10-080-CV), and 50% Neurobasal media (Gibco 21103049) supplemented with 0.5x N-2 (Gibco, 17502048), 0.5x B-27 (Gibco,17504044), 0.05% BSA (Gibco 15260-037), 100U/mL Penicillin and 0.1 mg/ml Streptomycin (Corning 30-002-CI), 1000 U/ml recombinant mouse LIF (Millipore ESG1106), 3 mM CHIR 99021 (STEMCELL Technologies 72054), 1 mM PD0325901 (STEMCELL Technologies 72184), 1.5×10^-4^ M 1-Thioglycerol (Sigma M6145) and 2 mM L-glutamine (Corning 25-005-CI). After thawing, cells were passaged once before differentiation and were not maintained in culture in 2i+LIF media for more than three passages. B-27 batches were tested to select only batches that maintained colonies with good morphology.

For differentiation, ES cells were dissociated with Accutase (Biolegend 423201) and plated in 100 x 15 mm Petri dishes (Falcon 351029) at a density of 40,000 cells/ml in 12.5ml serum-free differentiation media (SFD). SFD is composed of 75% IMDM (Corning 10-016-CM), 25% Ham’s F12 (Corning 10-080-CV), 0.5x N-2 (Gibco 17502048), 0.5x B-27 (Gibco 17504044), 0.05% BSA (Gibco 15260-037), 0.5mM Ascorbic Acid (Sigma A4544), 2mM Glutamine (Corning 25-005-CI), 0.45 mM 1-Thioglycerol (Sigma M6145), 100U/mL Penicillin and 0.1 mg/ml Streptomycin (Corning 30-002-CI). After two days, embryoid bodies (EBs) were dissociated with Accutase and resuspended at a density of 80,000 cells/ml in 4.5 ml SFD with Activin A (R&D Systems 338-AC) at the indicated concentration in Petri dishes (60 x 15 mm dishes, Falcon 351007). The differentiation efficiency was dependent on B-27 batches, so batches were tested to give higher than 50% DE progenitor at a concentration of 75 ng/ml of Activin A. For induction of Gata4 or Gata6, doxycycline (Sigma D9891) was added as a single dose of 30 ng/ml on day 3 (24 hours after Activin A supplementation). Stocks of doxycycline were maintained at −20 °C for no more than 3 months. Where indicated, WNT signaling activity was induced with CHIR 99021 (STEMCELL Technologies 72054) at the concentrations indicated. To inhibit Wnt signaling, 5 mM XAV-939 (Sigma-Aldrich X3004) was used.

For liver differentiation, day 5 EBs were dissociated with Accutase and plated at a ratio of 1:2 in SFD containing 50 ng/ml BMP4 (R&D Systems 314-BP), 10 ng/ml FGF2 (R&D Systems 233-FB), 50 ng/ml Activin A, and 10 ng/ml VEGF (R&D Systems 293-VE). After 24 hours, media was changed to SFD containing 50 ng/ml BMP4 (R&D systems), 10 ng/ml EGF (Peprotech 315-09), 10 ng/ml FGF2, 20 ng/ml HGF (Peprotech 100-39), 20 ng/ml TGFα (Peprotech 100-16A), 10 ng/ml VEGF and 10^−7^ M dexamethasone (Sigma D8893). Cells were maintained in this media for 2 days with daily media change.

For mid-hindgut differentiation, day 5 EBs were dissociated with Accutase and plated at a ratio of 1:2 in SFD supplemented with 10 ng/ml BMP4, 100 ng/ml FGF2, and 3 μM CHIR99021. Cells were maintained in the same media for 3 days with daily media changes.

### RNA isolation and RT-qPCR analysis

Total RNA was extracted using the RNeasy Plus Mini Kit (QIAGEN 74136), and 1 μg RNA was reverse transcribed using the SuperScript VILO cDNA Synthesis Kit (Invitrogen 11754250) following the manufacturer’s instructions. Quantitative PCR was carried out in triplicate with the LUNA universal qPCR Master mix (NEB M3003E) on a LightCycler 480 II instrument (Roche). Gene expression was normalized to endogenous Tbp. Primer sequences can be found in Table S1.

### RNA-seq

Total RNA was extracted from EBs 24h after treatment with Dox, CHIR, or DMSO with the RNeasy Plus Mini Kit (QIAGEN 74136) from three independent biological replicates per sample. Concentration was measured using a Nanodrop (Thermo Scientific), and RNA integrity was evaluated using a 2100 Bioanalyzer (Agilent Technologies). All processed samples had a greater than 9 RIN. Libraries were prepared using the NEBNext Ultra II Directional RNA Library Prep Kit for Illumina (New England Biolabs, USA), following the manufacturer’s protocol for use with the NEBNext Poly(A) mRNA Magnetic Isolation Module. Libraries were sequenced on the Illumina NovaSeq X Plus platform with paired-end 100 base pair reads at the Weill Cornell Medicine Genomics Resources Core Facility. Raw sequencing data in BCL format were converted to FASTQ files and demultiplexed using bcl2fastq v2.20 (Illumina). Reads from FASTQ files were aligned to the mouse genome (GRCm38.p6) and analyzed for differential expression using Basepair software with a Compare Gene Expression Levels using the DESeq2 pipeline (basepair.com). Differentially expressed genes (DEG) were defined as genes with a log_2_ fold change > 1 and p-adj < 0.05. Gene Set Enrichment Analysis (GSEA) was performed using GSEA 4.3.3 software (Broad Institute, Inc. and Regents of the University of California) ((Subramanian et al., 2005) on RNA-seq gene lists in which genes with a base mean of less than 50 counts. The comparison was performed using the mouse collections of the Molecular Signatures Database (MSigDB) mouse as a reference (Castanza et al., 2023) or custom gene lists compiled from literature as described in supplementary Table S2. Enrichment was considered significant for gene sets with a normalized enrichment score (NES) with a nominal p-value < 0.05 and a False Discovery Rate (FDR) q-value < 0.25. Heatmaps were generated from normalized counts from selected gene lists using SRplot (<u>bioinformatics.com.cn/srplot</u>) (Tang et al., 2023). Functional enrichment gene list analysis for transcription factors, pathways, and Gene Ontology (GO) terms was performed using the Enrichr web tool (https://maayanlab.cloud/Enrichr/) (Chen et al., 2013). For this analysis, the gene lists included only DEGs with a base mean of more than 100 counts.

### Flow cytometry

Live cells were used to detect cell surface markers. EBs were dissociated with Accutase, rinsed with FACS buffer (PBS containing 2mM EDTA and 1.5% IgG-free, Protease-Free BSA [Jackson ImmunoResearch 001-000-162]), and then incubated for 30 minutes on ice with primary antibodies diluted in FACS buffer. Subsequently, cells were rinsed twice with FACS buffer and stained for 5 mins at RT with SYTOX Blue Dead Cell Stain for flow cytometry (Invitrogen S34857) diluted 1:1000 in FACS buffer. Antibodies used for DE progenitors identification were CD117 (c-Kit) Monoclonal Antibody (2B8), PerCP-eFluor 710 conjugated (eBioscience 46-1171-82); CD184 (CXCR4) Monoclonal Antibody (2B11), APC conjugated (eBioscience 17-9991-82); and CD326 (EpCAM) Monoclonal Antibody (G8.8), PE-conjugated (eBioscience 12-5791-81). Antibodies against mesoderm markers were Dll1 Monoclonal Antibody (HMD1-3), PE-conjugated (Biolegend 128307); CD309 (VEGFR2, Flk-1) Monoclonal Antibody (Avas12), PE/Cyanine7 conjugated (Biolegend 136414); and CD140a (PDGFRA) Monoclonal Antibody (APA5), APC conjugated (eBioscience 17-1401-81).

For detection of Eomes and Brachyury, after EB dissociation, the cells were rinsed with PBS and then fixed with 2% PFA (Electron microscopy Sciences 15710) for 30 mins on ice. After centrifugation, cells were washed with PBS, resuspended in FACS buffer, and stored for up to a week at 4oC. Permeabilization and washes were done with the FoxP3 Transcription Factor Staining Buffer Set (eBioscience 00-5523-00) according to manufacturer instructions. After blocking with 5% FBS in permeabilization buffer for 15 minutes at RT, cells were incubated with PE-conjugated BRACHYURY polyclonal antibody (R&D Systems IC2085P), and Eomes Monoclonal antibody (W17001A), Alexa Fluor 647 conjugated (Biolegend 157703) for two hours at RT. After washing twice, cells were resuspended in FACS buffer and analyzed. All flow data was collected using an Attune NxT Flow Cytometer (ThermoFisher).

### Western Blotting

EBs were collected, washed with cold PBS, and lysed using RIPA buffer composed of 150 mM NaCl, 50 mM Tris-HCl pH 7.4, 1% NP-40, 0.5% sodium deoxycholate, 1x protease and phosphatase inhibitors (Pierce A32961) and 25 U/ml benzonase (Millipore 70664) on ice. After 10 minutes, SDS and EDTA were added to the lysate to a final concentration of 0.1% SDS and 2 mM EDTA, respectively. After an additional 20 min incubation on ice, the lysate was centrifuged at 15,000 × g for 15 min at 4°C. Aliquots were maintained at −80°C until analysis. The protein content was measured using the bicinchoninic acid protein assay kit (Pierce 23225). Proteins were then separated using NuPAGE 4 to 12% Bis-Tris Gels (Invitrogen NP0335BOX) before transferring to a PVDF membrane (Bio-Rad 162-0177). The membrane was blocked for 1 hour at room temperature using 5% IgG-free, Protease-Free BSA (Jackson ImmunoResearch 001-000-162) in 1x Tris-buffered saline plus 1% Tween 20 (TBST). After blocking, the blot was incubated with a primary antibody in 5% IgG-free BSA overnight at 4°C with gentle shaking. The next day, the membrane was washed 3 times using TBST. The secondary antibody was incubated for 1 hour at room temperature in TBST. Then, three washes were performed, and the membrane was developed with SuperSignal West Pico PLUS Chemiluminescent Substrate (Thermo Scientific 34577) and scanned using a chemiluminescence Licor’s C-DiGit imaging system. Primary antibodies used were GATA6 (Cell signaling technology 5851), GATA4 (Cell signaling technology 36966), SOX17 (R&D Systems AF1924), EOMES (Abcam ab23345), OCT3/4 (BD Transduction Laboratories 611202), and BRACHYURY (R&D Systems AF2085). Secondary antibodies used were Goat anti-rabbit IgG (H+L)-HRP Conjugate (BioRad 1706515), Goat Anti-Mouse IgG (H + L)-HRP Conjugate (BioRad 1706516), and Rabbit Anti-Sheep IgG (H+L)-HRP Conjugate (BioRad 1721017).

---

**Table S1.**
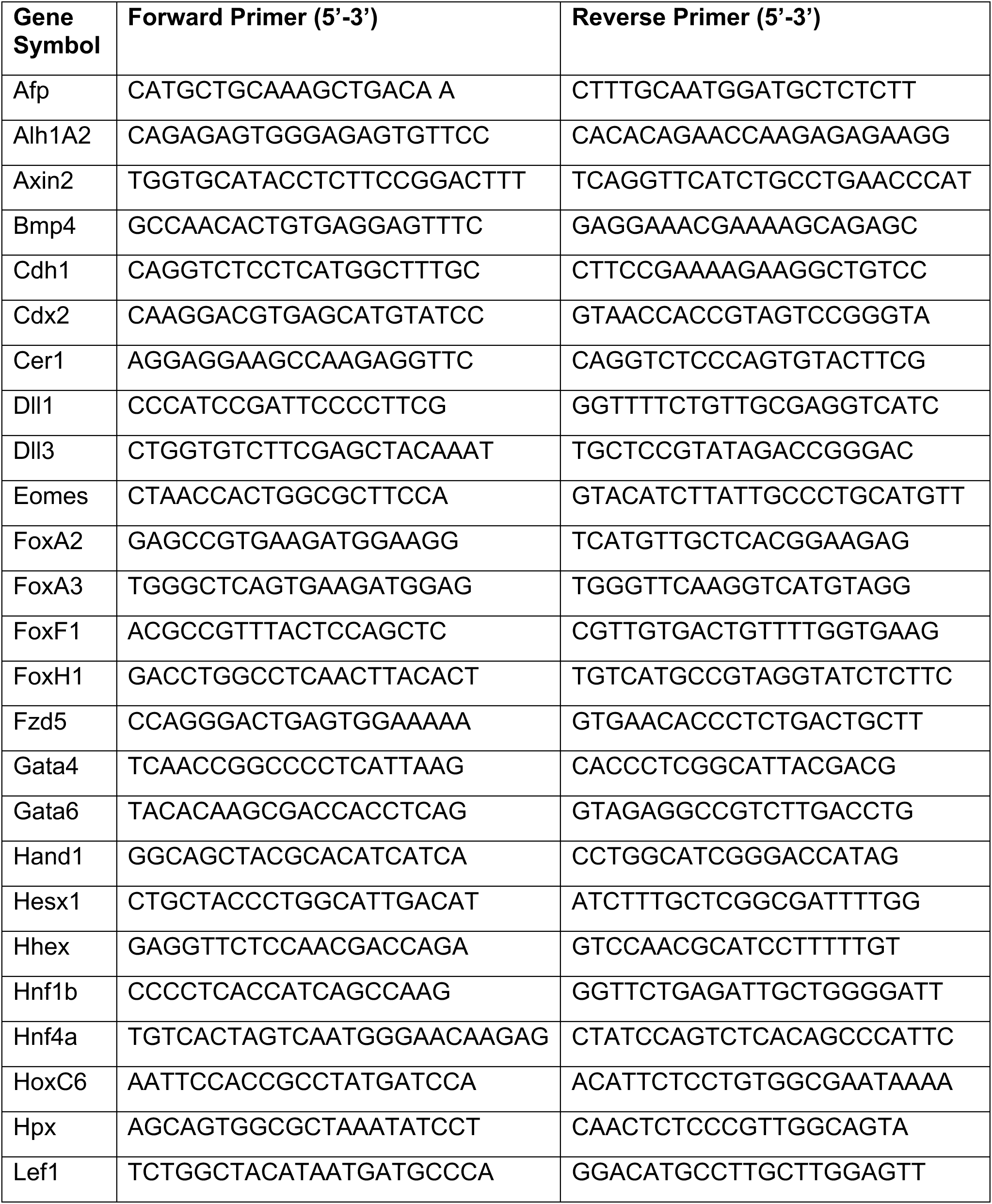
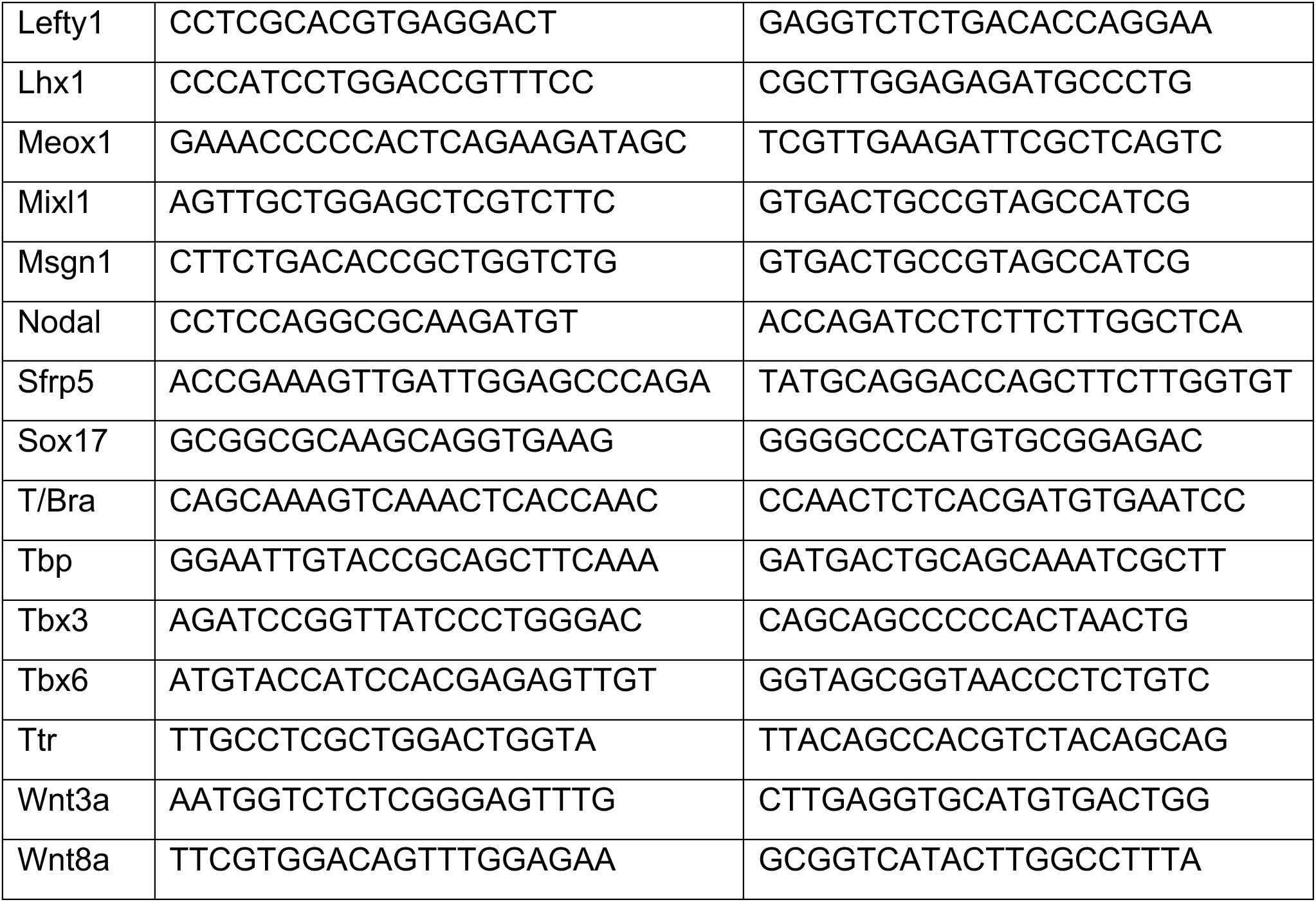
RT-qPCR primer sequences used in the study.

**Table S2.** Gene sets compiled from literature for GSEA analysis and heatmaps.

| **Reference** | **Source** | **Description** |
| --- | --- | --- |
| (Peng et al., 2019) | Supplementary Table 5 | Data analyzed corresponds to E7.0 and E7.5 stages. |
| (Ibarra-Soria et al., 2018) | Supplementary Table 2 | Sets analyzed include only genes with a Log2FC>1.5 for genes enriched in endoderm, and a Log2FC>1 for genes enriched in mesoderm and ectoderm. |
| (Pijuan-Sala et al., 2019) | Gene sets were extracted using the website https://marionilab.cruk.cam.ac.uk/MouseGastrulation2018/ accessed on February 15 2023 | To obtain the gene sets, definitive endoderm and nascent and mixed mesoderm at E7.25 and E7.5 were compared. |
| (Mariani et al., 2021) | Supplementary Table 2 | Gene sets analyzed include only those with more than 15 genes. |
| (Koh et al., 2016) | Data Citation 3. | Gene sets analyzed include lateral plate and paraxial mesoderm defined by the authors by evaluating expression of the genes after differentiation against expression in pluripotent cells. |
| (Kim et al., 2011) | Supplemental excel file | Gene sets were obtained by intersecting the list of high value targets with the lists of genes with SMAD binding peaks in hESCs and SMAD binding peaks in endoderm. Additionally, a gene set was defined for genes with a SMAD2 and 3 co-binding in endoderm. |
| (Chia et al., 2019) | Supplementary Table 1. | Gene set was obtained from the data of triple overlap of Gata6, Eomes and Smad2/3 ChIPseq data. |
| (Heslop et al., 2021) | Supplementary Table 3 | Gene sets analyzed were GATA6 dependent expression and accessibility, GATA6-dependent expression only, and GATA6-dependent accessibility only. |
| (Senft et al., 2018) | Supplementary Tables 1 and 2. | Gene sets were obtained by intersecting the RNAseq and ChIPseq data to obtain the sets of SMAD2/3 direct targets. |
| (Chiu et al., 2014) | Supplementary Table 3 | Gene sets were obtained for genes with SMAD2/3 and FOXH1 binding. |
| (Faial et al., 2015) | Supplementary Table S3 | Gene sets were prepared for genes positive or negatively regulated by Brachyury in response to Activin A and/or BMP. |
| (Tsaytler et al., 2023) | Supplementary table S5 | Gene sets were obtained from the columns of direct targets of SMAD1/5, SMAD2, EOMES, and BRACHYURY. |
| (Schüle et al., 2023) | Table S3 and S4 | Gene sets for high-confidence targets were obtained as described by the authors by intersecting downregulated genes in EOMES OR BRACHYURY knockout cells with genes rescued |
|  |  | in these KO cells after inducing the expression of EOMES or BRACHYURY. |
| (Harland et al., 2021) | Supplementary Table 6 and GSE140091 (RNAseq data) | Gene sets were obtained for Eomes direct targets by intersecting ChIPseq and RNAseq data. |
| (Zhang et al., 2013) | Supporting information Table S3 | Two sets of genes induced and repressed by WNT were extracted from Table S3b and S3c respectively. |
| (Estarás et al., 2015) | Supplementary Table S1 | A set was obtained from the column of genes that respond more strongly to the Wnt + Activin A combination. |
| (Koch et al., 2017) | Supplementary Table S2 | Genes bound by BRACHYURY and expression enriched in all Bra+ sorted populations. |

Gene lists collected from other species (human, frog etc.) were converted to mouse gene orthologs using Biomart at ensembl.org (Dyer et al., 2025). Similarly, gene lists collected from papers published before 2020 were updated using Biomart.

**Table S3.** Statistical analysis for Figure 6.

| Figure 6A. ANOVA test with multiple comparisons for Eomes expression (qPCR) |  |  |  |  |  |
| --- | --- | --- | --- | --- | --- |
| Tukey's multiple comparisons test | Mean Diff. | 95.00% CI of diff. | Below threshold? | Summary | Adjusted P Value |
| **D3.25** |  |  |  |  |  |
| -Dox DMSO vs. +Dox DMSO | -0.6604 | -0.9351 to -0.3856 | Yes | **** | <0.0001 |
| -Dox DMSO vs. -Dox CH 6uM | -0.3571 | -0.6318 to -0.08230 | Yes | ** | 0.0061 |
| -Dox DMSO vs. +Dox CH 6uM | -0.4135 | -0.6883 to -0.1387 | Yes | ** | 0.0012 |
| +Dox DMSO vs. -Dox CH 6uM | 0.3033 | 0.02853 to 0.5781 | Yes | * | 0.0252 |
| +Dox DMSO vs. +Dox CH 6uM | 0.2469 | -0.02790 to 0.5216 | No | ns | 0.0925 |
| -Dox CH 6uM vs. +Dox CH 6uM | -0.05643 | -0.3312 to 0.2183 | No | ns | 0.947 |
| **D3.5** |  |  |  |  |  |
| -Dox DMSO vs. +Dox DMSO | -1.979 | -2.253 to -1.704 | Yes | **** | <0.0001 |
| -Dox DMSO vs. -Dox CH 6uM | 0.1343 | -0.1405 to 0.4091 | No | ns | 0.5669 |
| -Dox DMSO vs. +Dox CH 6uM | -0.7986 | -1.073 to -0.5239 | Yes | **** | <0.0001 |
| +Dox DMSO vs. -Dox CH 6uM | 2.113 | 1.838 to 2.388 | Yes | **** | <0.0001 |
| +Dox DMSO vs. +Dox CH 6uM | 1.18 | 0.9052 to 1.455 | Yes | **** | <0.0001 |
| -Dox CH 6uM vs. +Dox CH 6uM | -0.9329 | -1.208 to -0.6582 | Yes | **** | <0.0001 |
| **D4** |  |  |  |  |  |
| -Dox DMSO vs. +Dox DMSO | -2.059 | -2.333 to -1.784 | Yes | **** | <0.0001 |
| -Dox DMSO vs. -Dox CH 6uM | 0.1063 | -0.1685 to 0.3811 | No | ns | 0.7329 |
| -Dox DMSO vs. +Dox CH 6uM | -1.804 | -2.079 to -1.530 | Yes | **** | <0.0001 |
| +Dox DMSO vs. -Dox CH 6uM | 2.165 | 1.890 to 2.440 | Yes | **** | <0.0001 |
| +Dox DMSO vs. +Dox CH 6uM | 0.2543 | -0.02043 to 0.5291 | No | ns | 0.0789 |
| -Dox CH 6uM vs. +Dox CH 6uM | -1.911 | -2.185 to -1.636 | Yes | **** | <0.0001 |

| Figure 6B. ANOVA test with multiple comparisons for T/Bra expression (qPCR) |  |  |  |  |  |
| --- | --- | --- | --- | --- | --- |
| Tukey's multiple comparisons test | Mean Diff. | 95.00% CI of diff. | Below threshold? | Summary | Adjusted P Value |
| **D3.25** |  |  |  |  |  |
| -Dox DMSO vs. +Dox DMSO | 0.4771 | -0.4301 to 1.384 | No | ns | 0.5058 |
| -Dox DMSO vs. -Dox CH 6uM | -3.87 | -4.777 to -2.962 | Yes | **** | <0.0001 |
| -Dox DMSO vs. +Dox CH 6uM | -1.593 | -2.500 to -0.6858 | Yes | *** | 0.0001 |
| +Dox DMSO vs. -Dox CH 6uM | -4.347 | -5.254 to -3.440 | Yes | **** | <0.0001 |
| +Dox DMSO vs. +Dox CH 6uM | -2.07 | -2.978 to -1.163 | Yes | **** | <0.0001 |
| -Dox CH 6uM vs. +Dox CH 6uM | 2.277 | 1.369 to 3.184 | Yes | **** | <0.0001 |
| **D3.5** |  |  |  |  |  |
| -Dox DMSO vs. +Dox DMSO | 0.7187 | -0.1886 to 1.626 | No | ns | 0.1651 |
| -Dox DMSO vs. -Dox CH 6uM | -9.958 | -10.86 to -9.050 | Yes | **** | <0.0001 |
| -Dox DMSO vs. +Dox CH 6uM | -2.127 | -3.034 to -1.219 | Yes | **** | <0.0001 |
| +Dox DMSO vs. -Dox CH 6uM | -10.68 | -11.58 to -9.769 | Yes | **** | <0.0001 |
| +Dox DMSO vs. +Dox CH 6uM | -2.845 | -3.753 to -1.938 | Yes | **** | <0.0001 |
| -Dox CH 6uM vs. +Dox CH 6uM | 7.831 | 6.924 to 8.738 | Yes | **** | <0.0001 |
| **D4** |  |  |  |  |  |
| -Dox DMSO vs. +Dox DMSO | 0.3849 | -0.5224 to 1.292 | No | ns | 0.6736 |
| -Dox DMSO vs. -Dox CH 6uM | -1.968 | -2.876 to -1.061 | Yes | **** | <0.0001 |
| -Dox DMSO vs. +Dox CH 6uM | -1.28 | -2.187 to -0.3728 | Yes | ** | 0.0026 |
| +Dox DMSO vs. -Dox CH 6uM | -2.353 | -3.261 to -1.446 | Yes | **** | <0.0001 |
| +Dox DMSO vs. +Dox CH 6uM | -1.665 | -2.572 to -0.7577 | Yes | **** | <0.0001 |
| -Dox CH 6uM vs. +Dox CH 6uM | 0.6883 | -0.2189 to 1.596 | No | ns | 0.1953 |

| Figure 6C. ANOVA test with multiple comparisons for EOMES+/BRACHYURY+ population |  |  |  |  |  |
| --- | --- | --- | --- | --- | --- |
| Tukey's multiple comparisons test | Mean Diff. | 95.00% CI of diff. | Below threshold? | Summary | Adjusted P Value |
| **D3** |  |  |  |  |  |
| -Dox DMSO vs. +Dox DMSO | 0 | -17.26 to 17.26 | No | ns | >0.9999 |
| -Dox DMSO vs. -Dox CH6 | 0 | -17.26 to 17.26 | No | ns | >0.9999 |
| -Dox DMSO vs. +Dox CH6 | 0 | -17.26 to 17.26 | No | ns | >0.9999 |
| +Dox DMSO vs. -Dox CH6 | 0 | -17.26 to 17.26 | No | ns | >0.9999 |
| +Dox DMSO vs. +Dox CH6 | 0 | -17.26 to 17.26 | No | ns | >0.9999 |
| -Dox CH6 vs. +Dox CH6 | 0 | -17.26 to 17.26 | No | ns | >0.9999 |
| **D3.5** |  |  |  |  |  |
| -Dox DMSO vs. +Dox DMSO | 1.144 | -16.12 to 18.40 | No | ns | 0.9978 |
| -Dox DMSO vs. -Dox CH6 | -22.53 | -39.79 to -5.274 | Yes | ** | 0.0073 |
| -Dox DMSO vs. +Dox CH6 | -33.98 | -51.24 to -16.72 | Yes | **** | <0.0001 |
| +Dox DMSO vs. -Dox CH6 | -23.68 | -40.94 to -6.417 | Yes | ** | 0.0047 |
| +Dox DMSO vs. +Dox CH6 | -35.12 | -52.38 to -17.86 | Yes | **** | <0.0001 |
| -Dox CH6 vs. +Dox CH6 | -11.45 | -28.71 to 5.815 | No | ns | 0.2847 |
| **D4** |  |  |  |  |  |
| -Dox DMSO vs. +Dox DMSO | -10.46 | -27.72 to 6.797 | No | ns | 0.3596 |
| -Dox DMSO vs. -Dox CH6 | -2.731 | -19.99 to 14.53 | No | ns | 0.9716 |
| -Dox DMSO vs. +Dox CH6 | -38.83 | -56.09 to -21.57 | Yes | **** | <0.0001 |
| +Dox DMSO vs. -Dox CH6 | 7.733 | -9.528 to 24.99 | No | ns | 0.6109 |
| +Dox DMSO vs. +Dox CH6 | -28.37 | -45.63 to -11.11 | Yes | *** | 0.0007 |
| -Dox CH6 vs. +Dox CH6 | -36.1 | -53.36 to -18.84 | Yes | **** | <0.0001 |

| Figure 6D. ANOVA test with multiple comparisons for EOMES+/BRACHYURY- population |  |  |  |  |  |
| --- | --- | --- | --- | --- | --- |
| Tukey's multiple comparisons test | Mean Diff. | 95.00% CI of diff. | Below threshold? | Summary | Adjusted P Value |
| **D3.5** |  |  |  |  |  |
| -Dox DMSO vs. +Dox DMSO | -41.12 | -51.01 to -31.24 | Yes | **** | <0.0001 |
| -Dox DMSO vs. -Dox CH6 | 7.29 | -2.595 to 17.18 | No | ns | 0.2036 |
| -Dox DMSO vs. +Dox CH6 | -0.208 | -10.09 to 9.677 | No | ns | >0.9999 |
| +Dox DMSO vs. -Dox CH6 | 48.41 | 38.53 to 58.30 | Yes | **** | <0.0001 |
| +Dox DMSO vs. +Dox CH6 | 40.91 | 31.03 to 50.80 | Yes | **** | <0.0001 |
| -Dox CH6 vs. +Dox CH6 | -7.498 | -17.38 to 2.387 | No | ns | 0.1841 |
| **D4** |  |  |  |  |  |
| -Dox DMSO vs. +Dox DMSO | -58.35 | -68.23 to -48.46 | Yes | **** | <0.0001 |
| -Dox DMSO vs. -Dox CH6 | 3.86 | -6.025 to 13.74 | No | ns | 0.7064 |
| -Dox DMSO vs. +Dox CH6 | -28.81 | -38.70 to -18.93 | Yes | **** | <0.0001 |
| +Dox DMSO vs. -Dox CH6 | 62.21 | 52.32 to 72.09 | Yes | **** | <0.0001 |
| +Dox DMSO vs. +Dox CH6 | 29.53 | 19.65 to 39.42 | Yes | **** | <0.0001 |
| -Dox CH6 vs. +Dox CH6 | -32.67 | -42.56 to -22.79 | Yes | **** | <0.0001 |

| Figure 6E. ANOVA test with multiple comparisons for EOMES-/BRACHYURY+ population |  |  |  |  |  |
| --- | --- | --- | --- | --- | --- |
| Tukey's multiple comparisons test | Mean Diff. | 95.00% CI of diff. | Below threshold? | Summary | Adjusted P Value |
| **D3.5** |  |  |  |  |  |
| -Dox DMSO vs. +Dox DMSO | 18.88 | 11.84 to 25.91 | Yes | **** | <0.0001 |
| -Dox DMSO vs. -Dox CH6 | -7.986 | -15.02 to -0.9527 | Yes | * | 0.0219 |
| -Dox DMSO vs. +Dox CH6 | 12.47 | 5.435 to 19.50 | Yes | *** | 0.0003 |
| +Dox DMSO vs. -Dox CH6 | -26.86 | -33.90 to -19.83 | Yes | **** | <0.0001 |
| +Dox DMSO vs. +Dox CH6 | -6.408 | -13.44 to 0.6256 | No | ns | 0.0832 |
| -Dox CH6 vs. +Dox CH6 | 20.45 | 13.42 to 27.49 | Yes | **** | <0.0001 |
| **D4** |  |  |  |  |  |
| -Dox DMSO vs. +Dox DMSO | 7.046 | 0.01237 to 14.08 | Yes | * | 0.0495 |
| -Dox DMSO vs. -Dox CH6 | -68.46 | -75.49 to -61.42 | Yes | **** | <0.0001 |
| -Dox DMSO vs. +Dox CH6 | 2.436 | -4.598 to 9.469 | No | ns | 0.7756 |
| +Dox DMSO vs. -Dox CH6 | -75.5 | -82.54 to -68.47 | Yes | **** | <0.0001 |
| +Dox DMSO vs. +Dox CH6 | -4.61 | -11.64 to 2.423 | No | ns | 0.2941 |
| -Dox CH6 vs. +Dox CH6 | 70.89 | 63.86 to 77.93 | Yes | **** | <0.0001 |

| EOMES+/BRACHYURY+ population |  |  |  |  |  |
| --- | --- | --- | --- | --- | --- |
| Tukey's multiple comparisons test | Mean Diff. | 95.00% CI of diff. | Below threshold? | Summary | Adjusted P Value |
| DMSO:-Dox A10 vs. DMSO:+Dox A10 | -10.46 | -27.64 to 6.718 | No | ns | 0.4493 |
| DMSO:-Dox A10 vs. XAV:-Dox A10 | 8.252 | -8.929 to 25.43 | No | ns | 0.7089 |
| DMSO:-Dox A10 vs. XAV:+Dox A10 | 0.2067 | -16.97 to 17.39 | No | ns | >0.9999 |
| DMSO:-Dox A10 vs. CH6:-Dox A10 | -2.731 | -19.91 to 14.45 | No | ns | 0.9991 |
| DMSO:-Dox A10 vs. CH6:+Dox A10 | -38.83 | -56.01 to -21.65 | Yes | **** | <0.0001 |
| DMSO:-Dox A10 vs. XCH:-Dox A10 | -0.578 | -17.76 to 16.60 | No | ns | >0.9999 |
| DMSO:-Dox A10 vs. XCH:+Dox A10 | -7.252 | -24.43 to 9.929 | No | ns | 0.8161 |
| DMSO:+Dox A10 vs. XAV:-Dox A10 | 18.72 | 1.534 to 35.90 | Yes | * | 0.0278 |
| DMSO:+Dox A10 vs. XAV:+Dox A10 | 10.67 | -6.511 to 27.85 | No | ns | 0.4267 |
| DMSO:+Dox A10 vs. CH6:-Dox A10 | 7.733 | -9.448 to 24.91 | No | ns | 0.7667 |
| DMSO:+Dox A10 vs. CH6:+Dox A10 | -28.37 | -45.55 to -11.19 | Yes | *** | 0.0007 |
| DMSO:+Dox A10 vs. XCH:-Dox A10 | 9.885 | -7.296 to 27.07 | No | ns | 0.5153 |
| DMSO:+Dox A10 vs. XCH:+Dox A10 | 3.211 | -13.97 to 20.39 | No | ns | 0.9974 |
| XAV:-Dox A10 vs. XAV:+Dox A10 | -8.045 | -25.23 to 9.136 | No | ns | 0.7324 |
| XAV:-Dox A10 vs. CH6:-Dox A10 | -10.98 | -28.16 to 6.198 | No | ns | 0.3935 |
| XAV:-Dox A10 vs. CH6:+Dox A10 | -47.08 | -64.26 to -29.90 | Yes | **** | <0.0001 |
| XAV:-Dox A10 vs. XCH:-Dox A10 | -8.83 | -26.01 to 8.351 | No | ns | 0.641 |
| XAV:-Dox A10 vs. XCH:+Dox A10 | -15.5 | -32.68 to 1.677 | No | ns | 0.0929 |
| XAV:+Dox A10 vs. CH6:-Dox A10 | -2.937 | -20.12 to 14.24 | No | ns | 0.9985 |
| XAV:+Dox A10 vs. CH6:+Dox A10 | -39.04 | -56.22 to -21.86 | Yes | **** | <0.0001 |
| XAV:+Dox A10 vs. XCH:-Dox A10 | -0.7847 | -17.97 to 16.40 | No | ns | >0.9999 |
| XAV:+Dox A10 vs. XCH:+Dox A10 | -7.459 | -24.64 to 9.722 | No | ns | 0.7955 |
| CH6:-Dox A10 vs. CH6:+Dox A10 | -36.1 | -53.28 to -18.92 | Yes | **** | <0.0001 |
| CH6:-Dox A10 vs. XCH:-Dox A10 | 2.153 | -15.03 to 19.33 | No | ns | 0.9998 |
| CH6:-Dox A10 vs. XCH:+Dox A10 | -4.521 | -21.70 to 12.66 | No | ns | 0.9805 |
| CH6:+Dox A10 vs. XCH:-Dox A10 | 38.25 | 21.07 to 55.43 | Yes | **** | <0.0001 |
| CH6:+Dox A10 vs. XCH:+Dox A10 | 31.58 | 14.40 to 48.76 | Yes | *** | 0.0002 |
| XCH:-Dox A10 vs. XCH:+Dox A10 | -6.674 | -23.85 to 10.51 | No | ns | 0.8684 |
| EOMES+/BRACHYURY- population |  |  |  |  |  |
| Tukey's multiple comparisons test | Mean Diff. | 95.00% CI of diff. | Below threshold? | Summary | Adjusted P Value |
| DMSO:-Dox A10 vs. DMSO:+Dox A10 | -58.35 | -72.12 to -44.58 | Yes | **** | <0.0001 |
| DMSO:-Dox A10 vs. XAV:-Dox A10 | -3.437 | -17.21 to 10.33 | No | ns | 0.9855 |
| DMSO:-Dox A10 vs. XAV:+Dox A10 | -61.07 | -74.84 to -47.30 | Yes | **** | <0.0001 |
| DMSO:-Dox A10 vs. CH6:-Dox A10 | 3.86 | -9.911 to 17.63 | No | ns | 0.9725 |
| DMSO:-Dox A10 vs. CH6:+Dox A10 | -28.81 | -42.58 to -15.04 | Yes | **** | <0.0001 |
| DMSO:-Dox A10 vs. XCH:-Dox A10 | 2.652 | -11.12 to 16.42 | No | ns | 0.9969 |
| DMSO:-Dox A10 vs. XCH:+Dox A10 | -62.39 | -76.16 to -48.62 | Yes | **** | <0.0001 |
| DMSO:+Dox A10 vs. XAV:-Dox A10 | 54.91 | 41.14 to 68.68 | Yes | **** | <0.0001 |
| DMSO:+Dox A10 vs. XAV:+Dox A10 | -2.726 | -16.50 to 11.04 | No | ns | 0.9963 |
| DMSO:+Dox A10 vs. CH6:-Dox A10 | 62.21 | 48.44 to 75.98 | Yes | **** | <0.0001 |
| DMSO:+Dox A10 vs. CH6:+Dox A10 | 29.53 | 15.76 to 43.30 | Yes | **** | <0.0001 |
| DMSO:+Dox A10 vs. XCH:-Dox A10 | 61 | 47.23 to 74.77 | Yes | **** | <0.0001 |
| DMSO:+Dox A10 vs. XCH:+Dox A10 | -4.044 | -17.81 to 9.726 | No | ns | 0.9649 |
| XAV:-Dox A10 vs. XAV:+Dox A10 | -57.64 | -71.41 to -43.87 | Yes | **** | <0.0001 |
| XAV:-Dox A10 vs. CH6:-Dox A10 | 7.296 | -6.474 to 21.07 | No | ns | 0.6083 |
| XAV:-Dox A10 vs. CH6:+Dox A10 | -25.38 | -39.15 to -11.61 | Yes | *** | 0.0002 |
| XAV:-Dox A10 vs. XCH:-Dox A10 | 6.089 | -7.682 to 19.86 | No | ns | 0.7812 |
| XAV:-Dox A10 vs. XCH:+Dox A10 | -58.95 | -72.72 to -45.18 | Yes | **** | <0.0001 |
| XAV:+Dox A10 vs. CH6:-Dox A10 | 64.93 | 51.16 to 78.70 | Yes | **** | <0.0001 |
| XAV:+Dox A10 vs. CH6:+Dox A10 | 32.26 | 18.49 to 46.03 | Yes | **** | <0.0001 |
| XAV:+Dox A10 vs. XCH:-Dox A10 | 63.73 | 49.95 to 77.50 | Yes | **** | <0.0001 |
| XAV:+Dox A10 vs. XCH:+Dox A10 | -1.318 | -15.09 to 12.45 | No | ns | >0.9999 |
| CH6:-Dox A10 vs. CH6:+Dox A10 | -32.67 | -46.44 to -18.90 | Yes | **** | <0.0001 |
| CH6:-Dox A10 vs. XCH:-Dox A10 | -1.208 | -14.98 to 12.56 | No | ns | >0.9999 |
| CH6:-Dox A10 vs. XCH:+Dox A10 | -66.25 | -80.02 to -52.48 | Yes | **** | <0.0001 |
| CH6:+Dox A10 vs. XCH:-Dox A10 | 31.47 | 17.70 to 45.24 | Yes | **** | <0.0001 |
| CH6:+Dox A10 vs. XCH:+Dox A10 | -33.58 | -47.35 to -19.81 | Yes | **** | <0.0001 |
| XCH:-Dox A10 vs. XCH:+Dox A10 | -65.04 | -78.81 to -51.27 | Yes | **** | <0.0001 |
| EOMES-/BRACHYURY+ population |  |  |  |  |  |
| Tukey's multiple comparisons test | Mean Diff. | 95.00% CI of diff. | Below threshold? | Summary | Adjusted P Value |
| DMSO:-Dox A10 vs. DMSO:+Dox A10 | 7.046 | -1.409 to 15.50 | No | ns | 0.141 |
| DMSO:-Dox A10 vs. XAV:-Dox A10 | 5.024 | -3.430 to 13.48 | No | ns | 0.4778 |
| DMSO:-Dox A10 vs. XAV:+Dox A10 | 7.192 | -1.263 to 15.65 | No | ns | 0.1273 |
| DMSO:-Dox A10 vs. CH6:-Dox A10 | -68.46 | -76.91 to -60.00 | Yes | **** | <0.0001 |
| DMSO:-Dox A10 vs. CH6:+Dox A10 | 2.436 | -6.019 to 10.89 | No | ns | 0.9682 |
| DMSO:-Dox A10 vs. XCH:-Dox A10 | -24.1 | -32.56 to -15.65 | Yes | **** | <0.0001 |
| DMSO:-Dox A10 vs. XCH:+Dox A10 | 6.933 | -1.521 to 15.39 | No | ns | 0.1525 |
| DMSO:+Dox A10 vs. XAV:-Dox A10 | -2.022 | -10.48 to 6.433 | No | ns | 0.9886 |
| DMSO:+Dox A10 vs. XAV:+Dox A10 | 0.146 | -8.309 to 8.601 | No | ns | >0.9999 |
| DMSO:+Dox A10 vs. CH6:-Dox A10 | -75.5 | -83.96 to -67.05 | Yes | **** | <0.0001 |
| DMSO:+Dox A10 vs. CH6:+Dox A10 | -4.61 | -13.06 to 3.844 | No | ns | 0.5765 |
| DMSO:+Dox A10 vs. XCH:-Dox A10 | -31.15 | -39.60 to -22.70 | Yes | **** | <0.0001 |
| DMSO:+Dox A10 vs. XCH:+Dox A10 | -0.1127 | -8.567 to 8.342 | No | ns | >0.9999 |
| XAV:-Dox A10 vs. XAV:+Dox A10 | 2.168 | -6.287 to 10.62 | No | ns | 0.9831 |
| XAV:-Dox A10 vs. CH6:-Dox A10 | -73.48 | -81.94 to -65.03 | Yes | **** | <0.0001 |
| XAV:-Dox A10 vs. CH6:+Dox A10 | -2.589 | -11.04 to 5.866 | No | ns | 0.9565 |
| XAV:-Dox A10 vs. XCH:-Dox A10 | -29.13 | -37.58 to -20.67 | Yes | **** | <0.0001 |
| XAV:-Dox A10 vs. XCH:+Dox A10 | 1.909 | -6.546 to 10.36 | No | ns | 0.9918 |
| XAV:+Dox A10 vs. CH6:-Dox A10 | -75.65 | -84.10 to -67.19 | Yes | **** | <0.0001 |
| XAV:+Dox A10 vs. CH6:+Dox A10 | -4.756 | -13.21 to 3.698 | No | ns | 0.5412 |
| XAV:+Dox A10 vs. XCH:-Dox A10 | -31.3 | -39.75 to -22.84 | Yes | **** | <0.0001 |
| XAV:+Dox A10 vs. XCH:+Dox A10 | -0.2587 | -8.713 to 8.196 | No | ns | >0.9999 |
| CH6:-Dox A10 vs. CH6:+Dox A10 | 70.89 | 62.44 to 79.35 | Yes | **** | <0.0001 |
| CH6:-Dox A10 vs. XCH:-Dox A10 | 44.35 | 35.90 to 52.81 | Yes | **** | <0.0001 |
| CH6:-Dox A10 vs. XCH:+Dox A10 | 75.39 | 66.94 to 83.85 | Yes | **** | <0.0001 |
| CH6:+Dox A10 vs. XCH:-Dox A10 | -26.54 | -34.99 to -18.08 | Yes | **** | <0.0001 |
| CH6:+Dox A10 vs. XCH:+Dox A10 | 4.498 | -3.957 to 12.95 | No | ns | 0.6039 |
| XCH:-Dox A10 vs. XCH:+Dox A10 | 31.04 | 22.58 to 39.49 | Yes | **** | <0.0001 |

| Tukey's multiple comparisons test | Predicted (LS) mean diff. | 95.00% CI of diff. | Below threshold? | Summary | Adjusted P Value |
| --- | --- | --- | --- | --- | --- |
| **DMSO** |  |  |  |  |  |
| WT vs. G6KO | 25.07 | 16.30 to 33.83 | Yes | *** | 0.0001 |
| **CH0.5** |  |  |  |  |  |
| WT vs. G6KO | 24.51 | 15.74 to 33.27 | Yes | *** | 0.0001 |
| **CH1** |  |  |  |  |  |
| WT vs. G6KO | 20.71 | 11.95 to 29.48 | Yes | *** | 0.0005 |
| **CH3** |  |  |  |  |  |
| WT vs. G6KO | 19.53 | 10.77 to 28.30 | Yes | *** | 0.0007 |
| **CH6** |  |  |  |  |  |
| WT vs. G6KO | 6.116 | -4.620 to 16.85 | No | ns | 0.2297 |
| **WT** |  |  |  |  |  |
| DMSO vs. CH0.5 | -3.106 | -30.64 to 24.42 | No | ns | 0.9968 |
| DMSO vs. CH1 | -1.449 | -28.98 to 26.08 | No | ns | 0.9998 |
| DMSO vs. CH3 | 10.68 | -16.85 to 38.21 | No | ns | 0.7663 |
| DMSO vs. CH6 | 31.19 | 0.4083 to 61.97 | Yes | * | 0.0462 |
| CH0.5 vs. CH1 | 1.657 | -25.87 to 29.19 | No | ns | 0.9997 |
| CH0.5 vs. CH3 | 13.78 | -13.75 to 41.31 | No | ns | 0.5672 |
| CH0.5 vs. CH6 | 34.29 | 3.514 to 65.07 | Yes | * | 0.0248 |
| CH1 vs. CH3 | 12.13 | -15.40 to 39.65 | No | ns | 0.6759 |
| CH1 vs. CH6 | 32.64 | 1.857 to 63.42 | Yes | * | 0.0347 |
| CH3 vs. CH6 | 20.51 | -10.27 to 51.29 | No | ns | 0.2985 |
| **G6KO** |  |  |  |  |  |
| DMSO vs. CH0.5 | -3.667 | -31.20 to 23.86 | No | ns | 0.994 |
| DMSO vs. CH1 | -5.804 | -33.33 to 21.73 | No | ns | 0.9668 |
| DMSO vs. CH3 | 5.139 | -22.39 to 32.67 | No | ns | 0.9786 |
| DMSO vs. CH6 | 12.23 | -18.54 to 43.01 | No | ns | 0.7503 |
| CH0.5 vs. CH1 | -2.136 | -29.67 to 25.39 | No | ns | 0.9993 |
| CH0.5 vs. CH3 | 8.807 | -18.72 to 36.34 | No | ns | 0.8662 |
| CH0.5 vs. CH6 | 15.9 | -14.88 to 46.68 | No | ns | 0.5383 |
| CH1 vs. CH3 | 10.94 | -16.59 to 38.47 | No | ns | 0.7503 |
| CH1 vs. CH6 | 18.04 | -12.74 to 48.82 | No | ns | 0.4185 |
| CH3 vs. CH6 | 7.095 | -23.68 to 37.87 | No | ns | 0.9545 |

| Tukey's multiple comparisons test | Mean Diff. | 95.00% CI of diff. | Below threshold? | Summary | Adjusted P Value |
| --- | --- | --- | --- | --- | --- |
| **DMSO** |  |  |  |  |  |
| WT vs. G6KO | -10.92 | -20.51 to -1.338 | Yes | * | 0.0275 |
| **CH0.5** |  |  |  |  |  |
| WT vs. G6KO | -18.24 | -27.82 to -8.651 | Yes | *** | 0.0008 |
| **CH1** |  |  |  |  |  |
| WT vs. G6KO | -17.6 | -27.19 to -8.019 | Yes | ** | 0.001 |
| **CH3** |  |  |  |  |  |
| WT vs. G6KO | -15.65 | -25.23 to -6.063 | Yes | ** | 0.0028 |
| **CH6** |  |  |  |  |  |
| WT vs. G6KO | -9.556 | -19.14 to 0.02785 | No | ns | 0.0506 |
| **WT** |  |  |  |  |  |
| DMSO vs. CH0.5 | -5.914 | -19.66 to 7.834 | No | ns | 0.7017 |
| DMSO vs. CH1 | -15.15 | -28.90 to -1.405 | Yes | * | 0.0264 |
| DMSO vs. CH3 | -58.8 | -72.55 to -45.05 | Yes | **** | <0.0001 |
| DMSO vs. CH6 | -73.24 | -86.99 to -59.50 | Yes | **** | <0.0001 |
| CH0.5 vs. CH1 | -9.24 | -22.99 to 4.509 | No | ns | 0.2967 |
| CH0.5 vs. CH3 | -52.88 | -66.63 to -39.13 | Yes | **** | <0.0001 |
| CH0.5 vs. CH6 | -67.33 | -81.08 to -53.58 | Yes | **** | <0.0001 |
| CH1 vs. CH3 | -43.64 | -57.39 to -29.89 | Yes | **** | <0.0001 |
| CH1 vs. CH6 | -58.09 | -71.84 to -44.34 | Yes | **** | <0.0001 |
| CH3 vs. CH6 | -14.45 | -28.20 to -0.6982 | Yes | * | 0.0365 |
| **G6KO** |  |  |  |  |  |
| DMSO vs. CH0.5 | -13.23 | -26.98 to 0.5211 | No | ns | 0.0629 |
| DMSO vs. CH1 | -21.83 | -35.58 to -8.086 | Yes | ** | 0.001 |
| DMSO vs. CH3 | -63.52 | -77.27 to -49.77 | Yes | **** | <0.0001 |
| DMSO vs. CH6 | -71.88 | -85.63 to -58.13 | Yes | **** | <0.0001 |
| CH0.5 vs. CH1 | -8.607 | -22.36 to 5.142 | No | ns | 0.3626 |
| CH0.5 vs. CH3 | -50.29 | -64.04 to -36.55 | Yes | **** | <0.0001 |
| CH0.5 vs. CH6 | -58.65 | -72.40 to -44.90 | Yes | **** | <0.0001 |
| CH1 vs. CH3 | -41.69 | -55.44 to -27.94 | Yes | **** | <0.0001 |
| CH1 vs. CH6 | -50.04 | -63.79 to -36.29 | Yes | **** | <0.0001 |
| CH3 vs. CH6 | -8.356 | -22.10 to 5.393 | No | ns | 0.3909 |

